# Predicting animal movement across the species range for landscape-scale applications

**DOI:** 10.1101/2025.06.12.658668

**Authors:** Kayleigh Chalkowski, Alexandra Sack, George Wittemyer, Ryan S. Miller, Raoul K. Boughton, William S. Raymond, Nathan P. Snow, Jim C. Beasley, Stephen S. Ditchkoff, Tyler S. Evans, Steven M. Gray, Jesse S. Lewis, Mitchell A. Parsons, Kurt C. VerCauteren, Stephen L. Webb, Julie K. Young, Kim M. Pepin

## Abstract

**Background:** Understanding animal movement across broad ecological contexts is essential for guiding management decisions in natural resource and animal disease management. However, most movement studies collect data over small spatial extents, which challenges using the data for landscape-scale management applications.

**Methods:** We collated relocation data from 564 wild pigs spanning 25 studies across 10 states in the United States. We derived net and daily displacement distributions for individuals in each of 4 seasons. We fit gamma distributions to displacement distributions and derived mean and dispersion parameters as metrics of displacement magnitude and variation. We modeled mean and dispersion variables using gradient-boosted regression, including landscape and weather predictors, and predicted daily and net displacement distributions across the species range. We used integrated step selection analysis to estimate habitat selection for wild pig groups sharing a common composition of habitat availability across the studied populations.

**Results:** Female average (**±** 95% confidence interval) net displacement means varied by season, peaking in Jul-Sept at 1,152 **±** 140 m along with daily displacement mean distances of 650 **±** 41 m. Average net displacement means for males peaked in Oct.-Dec. at 1,651 **±** 160 m with daily displacement mean distances of 1,092 **±** 95 m. Extreme net displacement values ranged up to 9,649 m and 4,080 m for male and female, respectively, and extreme daily displacement distribution means of 6,616 m for males, and 3,855 m for females. Almost half of the individuals included a similar composition of habitat availability, regardless of study site location. Herbaceous wetland was the most consistently favored land type and was available for more than 70% of all individual pigs across all studies. Preference for other land types depended strongly on habitat availability.

**Conclusions:** By comparing movement and habitat selection across a large geographic extent, our work provides new insights about the environmental drivers and limits of wild pig movement behavior. Our workflow digests variation wild pig movement distributions and habitat selection into parameters that can be used to predict movement on heterogeneous landscapes for downstream applications such as invasive species elimination campaigns or disease management.

## Introduction

Animal movement underpins numerous ecological processes including disease spread [1–3], invasive species spread [4], and population persistence [5]. As such, characterizing species movement behavior is fundamental to wildlife conservation and management efforts [6]. For example, infrastructure development projects worldwide require predictions of how proposed development changes might alter animal movements or population dynamics [7]. More generally, conservation land management decisions are improved by consideration of how animals move through the landscape to best support healthy populations [8]. Wildlife disease can be managed most effectively when knowledge of animal movement through landscapes is leveraged towards identifying optimal spatial distribution of disease control [9]. However, management decisions involving these processes rely on understanding animal movement across large geographic extents, including regions where animal relocation data have not been collected [10]. Prediction of animal movement out of sample is typically challenging and frequently limits the use of these data to inform management activities [11].

The movement ecology field is grounded in understanding movement decisions of individual animals from relocation data [12–14]. As global positioning system (GPS) technology has improved, so has the emphasis on understanding animal movement behavior at finer spatial scales [15]. For example, path-level analyses aimed at characterizing the step length or turning angles along an individual’s path of movement (e.g., [16]), path segmentation methods to understand an individual’s behavioral state along the path of movement (e.g., [17,18] or methods that relate fine-scaled movement behavior to habitat features [19,20]. These advances in understanding animal movement mechanisms on heterogeneous landscapes have laid the foundation for predicting animal movement in new areas from mechanistic principles [10,11,21–23]. In particular, deep learning approaches can predict movement trajectories and estimate step probabilities with very high accuracy while allowing for nonlinear relationships with environmental predictors -- offering greater flexibility in handling complex non-linear relationships across space [11]. However, predicting individual movement trajectories at a high resolution comes with substantial computational costs [11] that may be infeasible for application to management issues across large geographic extents.

For some applications, predicting movement behavior at coarse scales is sufficient for landscape-scale management decisions where interventions occur broadly, such as planning disease control [21,24], conservation land planning [25], or efficient allocation of resources for eliminating invasive species [26,27]. For applications focused on inter-individual or inter-specific interactions, detailed analysis can provide valuable insights [28] but often is computationally difficult to implement over large sample sizes and spatial scales. Individual-level summaries of animal movement metrics (e.g., distance, site fidelity) could be useful for representing animal movement behavior and its heterogeneity, while identifying governing mechanisms of movement in different ecological contexts. Generating a predictive understanding of individual-level variation in animal movement across environmental contexts, in turn, allows for predicting movement behavior across the species range, providing data and insight for ecological process models (e.g., disease dynamics or invasive species spread) that inform management decisions at relevant spatial and temporal scales [10].

Wild pigs are a globally distributed species that cause immense damage to agriculture, natural resources, and personal property [29] and threaten animal and human health [30,31]. In the United States (U.S.) wild pigs are an invasive species originating from a mixed ancestry of domestic pig and wild boar [32]. Their current distribution spans the southern half of the U.S. [33] with occasional northern populations that emerge from translocation events [34]. A national management program led by the United States Department of Agriculture Animal and Plant Health Inspection Service Wildlife Services since 2014 has been effective at curtailing historical rates of population expansion [33,35–37] but long established populations continue to cause damage and disease risks in some areas [38]. Animal movement studies across the range of the wild pig distribution in the U.S. have revealed that movement behavior can differ dramatically across geographic contexts, due to factors such as meteorological variables, habitat, season, agriculture, and individual-level characteristics [39,40] that have downstream effects for planning management of animal diseases [24] and mitigating expansion of the species [36]. Wild pig movement analyses have provided important insights for understanding how wild pigs use heterogeneous landscapes in local areas [41] and knowledge and methods for predicting habitat-driven movements of individuals. These advancements provide data and methods that can be leveraged further towards landscape-scale management challenges such as rapid control of translocations to new areas or emergency management of foreign animal diseases [24,38].

The objective of our study was to derive individual-level metrics of wild pig movement at a spatio-temporal scale that can be integrated directly for prediction of population-level ecological processes that inform landscape-scale management decisions. We were especially interested in providing movement metrics for emergency management planning of disease (e.g. [24]). A key need in preparedness for animal disease emergencies is prediction about the rate of spatial spread of disease in different geographic areas. In wildlife, disease spread depends on animal movement behavior [2,3] that may vary dramatically across ecological contexts [10].

Thus, characterizing the variation in fundamental animal movement features [13] across the species range is an essential ingredient for guiding disease management preparedness. The key movement features we quantified were: movement distance to new areas on the landscape over time relative to an initial location of residency (net displacement; a measure of site fidelity), distance between day-to-day shifts in where individuals reside (daily displacement; a measure of movement shifts about a home range), and where to reside (habitat selection). Our analysis provides insight for mechanistic understanding and prediction of wild pig movement across large geographic extents where the habitat composition changes dramatically. More broadly, our application provides an informative workflow for estimating individual-level movement parameters across a range of environmental contexts that can be used to parameterize ecological process models that depend on realistic rules of animal movement. We discuss the utility of our approach for providing reasonable approximations of individual-level movement distributions across the species range for terrestrial species with habitat-driven movement.

## Methods

### Overview

All analyses were conducted in R version 4.2.0 [42]. We analyzed relocation data from 564 individual wild pigs across 25 study sites across 10 states (Figure 1). We developed separate analysis workflows for estimating net and daily displacement distance (Figure 2) and habitat selection. We summarized variation in individual wild pig displacement distances by fitting each individual’s distribution of displacement distances to a gamma distribution. This allows characterization of an individual’s displacement distributions (net and daily) using only two parameters per distribution. We then leveraged the predictive power of boosted regression [43] to quantify environmental predictors of wild pig displacement distributions for watersheds where wild pigs were sampled. We used this fitted model to predict wild pig displacement distribution features across the entire species’ range in the U.S., which enables extraction of the full wild pig displacement distributions across the range of prediction.

**Figure 1.**
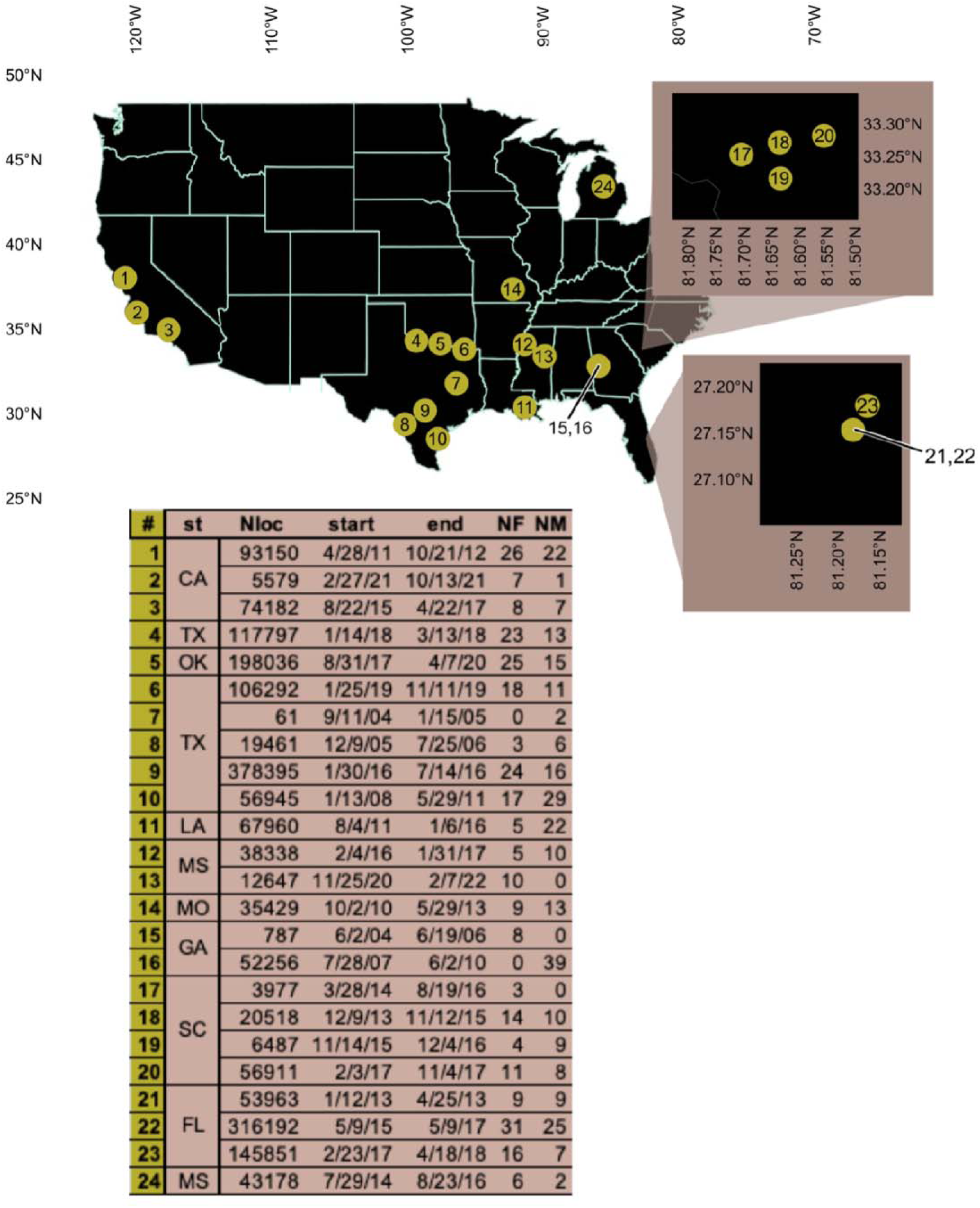
Locations of U.S. wild pig (*Sus scrofa*) GPS collaring studies included in this analysis with total numbers of wild pigs per study, total numbers of females (NF), males (NM) with number of geolocations (Nloc) and start and end dates of geolocation data use in analyses.

**Figure 2.**
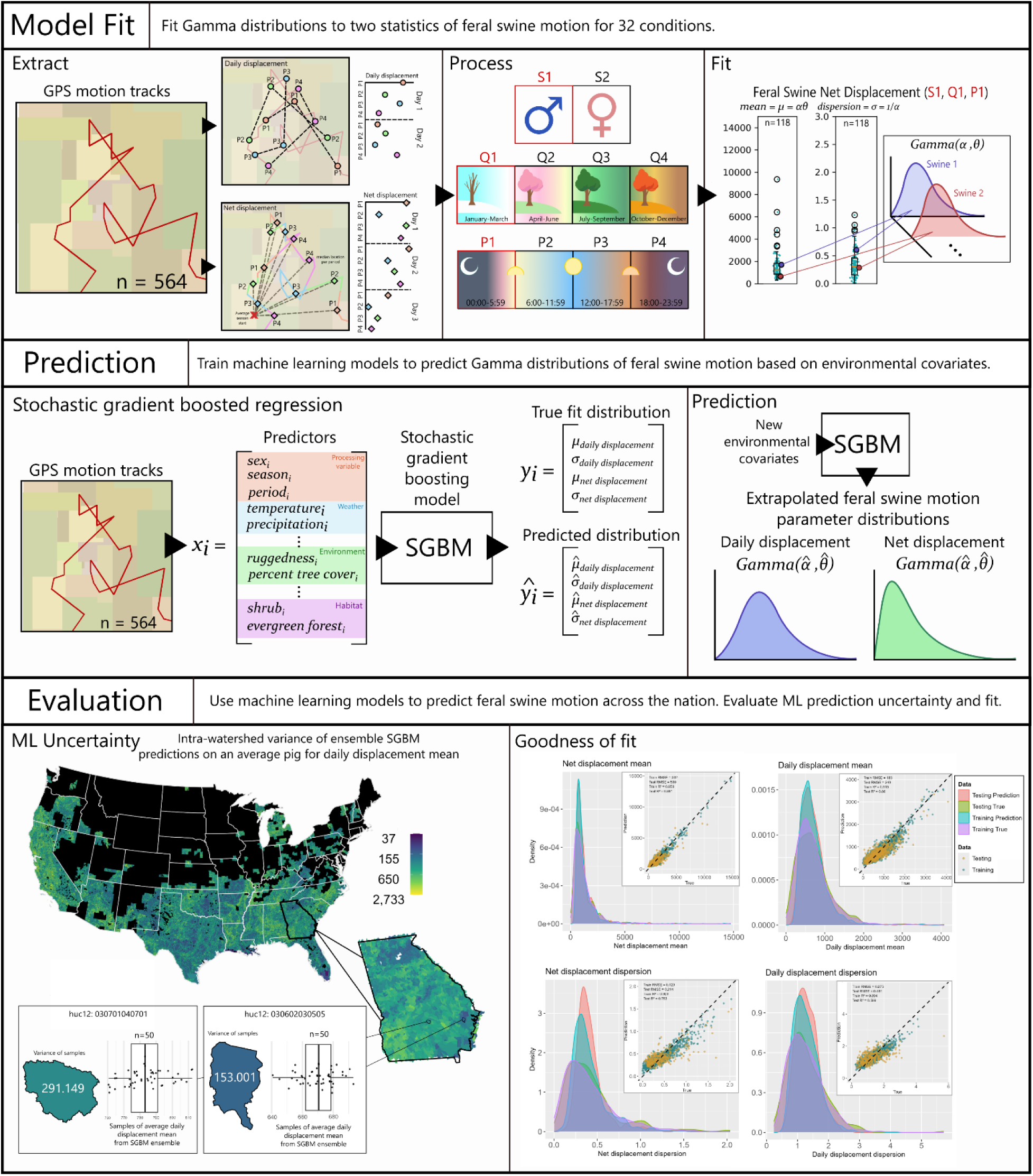
Conceptual workflow for modeling and predicting wild pig net and daily displacement. Observed net and daily displacement distances from wild pig relocation data (*i*, *N*=564 wild pigs) were fit with gamma distributions to estimate shape and scale parameters. We then derived the mean (μ*_i_* = shape x scale) and dispersion (σ*_i_* = 1/shape) features of these displacement distributions for each individual wild pig for different seasons and periods of the day, providing measures of central tendency and skew. These four responses (mean and dispersion for each of net and daily displacement distribution) were then modeled separately as functions of environmental covariates (*x_i_*) using stochastic gradient boosted (SGB) regression. For predicting movements of individuals in locations *j*, replicate SGB ensembles were used to predict movement features (μ_ij_, σ_ij_) from environmental covariates for a representative individual in location *j*. The predicted movement features were then used to derive shape and scale parameters for a representative individual in a given location to simulate the distributions of net and daily displacement behavior. Uncertainty in out-of-sample model predictions was estimated using the coefficient of variation of individual model predictions from the ensemble of models in each watershed, providing a measure of precision of watershed-level predictions. Finally, to evaluate goodness-of-fit of the models, we compared the distributions of predicted and observed response variables for each season and estimated the squared correlation coefficient among observed and predicted response data.

Finally, we estimated habitat selection coefficients to quantify how land cover availability may determine habitat selection. A key challenge with estimating habitat selection across broad spatial scales is that available habitat can vary dramatically. We addressed this challenge by subsetting individuals into groups that shared a common set of available landcovers in their home ranges and then estimated habitat selection for each subset of individuals. We then took the average and variation of each coefficient in the group to summarize habitat selection profiles for each grouping of habitat availability.

### Processing geolocation data

Data were filtered for geolocations with more than 32 km/hr (20 mph) between points using the trip package [44] and pigs with GPS-tracking for at least five calendar days. Temporal resolution of GPS fixes varied substantially across study sites, which was accounted for by resampling geolocations at a fix rate of every six hours (4 locations per day per individual) to standardized fix rates across study site, using the amt package [45]. Keeping 4 locations per day per individual allowed us to quantify daily movement metrics during 4 different wild pig activity periods of the day. The four daily partitions were 0:00-5:59, 6:00-11:59, 12:00-17:59, and 18:00-23:59 to account for known differences in wild pig activity patterns at different times of the day [46]. We subset relocation data into four equally long seasons: January-March; April-June; July-September; and October-December, to compare changes in movement metrics throughout the calendar year.

We derived two movement metrics from the relocation data for each daily period and season: <u>net displacement</u> (calculated for each day during the season) and <u>daily displacement</u>. Net displacement was defined as the Euclidean distance between the wild pig’s starting point of relocation data (the average location for the first seven days of each season during each period of the day) and median daily location within each of the four time periods (i.e., generating four net displacement distances per day) using the sf library [47]. Daily displacement was defined as the distance between a randomly sampled geolocation within the same 6-hour period at 24-hour intervals, allowing a sampling tolerance range of fixes that were 24**±**3 hours apart between days during similar periods of the day (Figure 2).

### Fitting displacement distributions for individuals

We fit gamma distributions to net and daily displacement distributions (separately) for each collared pig/day period/season combination using the fitdistrplus package [48]. We chose a unimodal gamma distribution because wild pig movements are generally short with some longer range movements [40] that we hypothesized could be accounted for through the skew in a gamma distribution. The gamma distribution is characterized by scale and shape parameters, which describe the magnitude and skew (shape) of the distribution and its spread (scale). We estimated the shape and scale parameters of each individual’s distribution of displacements (separately). Then, we chose two distinct features of these distributions as response variables to quantify effects of environmental covariates - ‘mean’ (shape x scale) and ‘dispersion’ (1/shape). This enables an understanding of how environmental covariates predict the average and skew of movement distances and prediction of two distribution features that can be used to simulate movement behavior anywhere the estimates are made. This led to 4 response variables in our modeling: net displacement means, net displacement dispersion, daily displacement means, and daily displacement dispersion.

### Estimating effects of environmental predictors on movement distance distributions

1. *<u>Modelling technique</u>*. We used gradient-boosted regression modeling to estimate environmental covariate effects on displacement response variables of individual pigs, due to its strength with handling complex, nonlinear ecological relationships [49]. We developed a separate model for each response variable. All boosted regression models were conducted using the *dismo* library [50] with a Poisson loss function for distribution means and a least squares loss function for distribution dispersion. We used the full dataset of 564 pigs for modeling effects of environmental covariates on response variables. However, since variance is sensitive to sample size, we used a subset of the data to model the dispersion responses that only included pigs with at least 30 measurements after geo-location data pre-processing. Prior to fitting the response data to environmental predictors, we evaluated spatial autocorrelation of the response variables between study sites using Moran’s I calculated with the ape library [51]. We did not find significant spatial autocorrelation between sites and thus did not include this effect in the models.
2. *<u>Predictor data processing</u>*. Environmental predictors were chosen based on variables known to be important for wild pig ecology from previous studies (e.g., [40]). Environmental predictors were spatially and temporally aggregated on a per-individual basis to match the scale of the response data (Table S1). The mean and variance estimates of each environmental predictor across the spatial and temporal scale of the relocation data for each individual were used as predictors. We estimated the environmental predictor variables for each watershed across the U.S. wild pig distribution for four separate seasons (Jan-Mar, Apr-Jun, Jul-Sep, Oct-Dec) to account for known effects of seasonality on wild pig movement [40] For variables that changed temporally at scales smaller than one year (i.e., temperature, precipitation, daylight range), we used the value for the middle month of each month range (i.e., 15 Feb, 15 May, 15 Aug, 15 Nov).
3. *<u>Model fitting and selection.</u>* Initial hyper parameter exploration was conducted on random and spatially-blocked [52] *n* x *k*-fold nested CV methods for optimizing the following hyperparameters:number of trees, learning rate, tree complexity, and bag fraction..We first optimized learning rate across a sequence (0.3, 0.1, 0.05, 0.01, 0.005, 0.001, 0.0005, 0.0003, 0.0001) while keeping other hyperparameters constant (tree complexity: 5; bag fraction: 0.5), and selecting the optimum number of trees from a sequence (from 100 to 8000, intervals of 100) for each learning rate. Then, we tuned the tree-specific parameters for the selected learning rate (tree complexity: 1, 3, 5, 7, 9, 10, 12; bag fraction: 0.2, 0.5, 0.6, 0.8, 0.9). From this previous grid exploration, a manual selection from the grid to minimize out-of-sample RMSE was performed for a simple 80:20 split of the data. Manually selected hyperparameters are listed in Table S2.
4. *<u>Prediction</u>*. We extracted predictor data for all watersheds across the wild pig range and aggregated predictors spatially within watersheds (habitat/landscape predictors), and temporally by our four seasonal periods (weather predictors). For continuous variables, we took the average and variance of all values within each watershed-season subset (where the sample size of values within watersheds was dependent on the spatial resolution of each variable and the size of the watershed; Table S1). For categorical variables (land class), we calculated the mean and variance of the proportion of each category in each watershed. Using the selected models (as above; Table S2), we predicted response variables for watersheds across the U.S. using a stochastic gradient boosted (SGB) ensemble [54,55]. We created a stochastic gradient-boosted (SGB) ensemble comprised of 100 (*M*) gradient-boosted decision trees (GBDT), each predicting net and daily displacement means and dispersion according to the environmental parameters. We used the mean of predictions from the ensemble of models from each watershed by season combination as the final prediction. For each watershed by season combination, we used the predicted mean and dispersion estimates to derive the shape and scale parameters for the expected gamma distribution of wild pig displacement distributions. Using the shape and scale parameters for net and daily displacement distributions, we simulated 1,000 random draws to represent the predicted displacement distributions for each watershed.
5. *<u>Evaluation of model fit and uncertainty</u>*. We estimated the epistemic uncertainty of our models using the variance of the resulting SGB predictions [55,56], reflected by the disagreement of the ensemble model predictions [55,57,58]. This provided a measure of precision for in- and out-of-sample predictions in each watershed across the wild pig range. We normalized the relative variances between watershed level predictions between 0 and 1 via min-max normalization for visualization purposes (Figure S1). We estimated absolute goodness of fit for the fitted models (in-sample predictions) using the squared correlation coefficient of the observed and predicted response variables for both training data and withheld testing data (80:20 split respectively) (Figure S2).

### Estimating habitat selection profiles

Given context-dependence of habitat selection [59], we assigned individuals into 12 groups based on the composition of habitat availability of four key land cover types: forest, cropland, wetland, and urban. The four land cover types have all been supported by previous research to be important to wild pig movement or habitat selection [60–66]. The largest habitat availability group contained 219 individuals. Each study site had at least one individual in the largest group and this group contained the four landcover types used for grouping.

We used integrated step selection analysis (iSSA), which uses a conditional logistic regression model [20]. The iSSA model estimates movement and resource selection parameters and allows inference about habitat selection within a mechanistic movement model. The data were resampled to every six hours and daily steps (24 hours with 3 hour window) were calculated four times a day. Individuals without a minimum of ten consecutive days were removed from the model. For the iSSA model, ten random steps were calculated using the *amt* package [45] drawn from the full distribution of steps and turn angles across all years and study sites. Each individual was run in a separate conditional regression model to examine habitat selection using deviation coding to compare the mean for each land cover type to the overall mean of land types. Only land cover types that each group of wild pigs visited at least once was included in each model. The average was then taken of all estimates that were significant for each habitat availability subset. All available land cover types from the National Land Cover Database (NLCD) were included, with the four levels of developed land combined into a single category of developed land. Our model was tested for its sensitivity of inclusion of step length as an interaction term, considering significant interactions, biological relevance, and AIC. Occasional long, regular movements were investigated for individuals to verify they were not errors.

To understand data-driven uncertainty in our top model, we also conducted a leave-one-out sensitivity analysis. In our leave-one-out analysis within each habitat availability subset, one study was removed at a time, the model was rerun, and the mean squared error was calculated for each group and the full model. The standard error was calculated for the estimates for each land type and each availability subset for every subset with more than one study from the leave-one out analysis. To further characterize uncertainty in our findings, we compared the correlation of these standard error results based on the number of studies, individuals, and total observations per availability subset using Pearson’s correlation.

## Results

### Trends in wild pig movement distances

Net and daily displacement means in males tended to be higher and more variable during Jul.-Dec. relative to the first half of the year (Figure 3, top). In general, net and daily displacement means of females were lower than males and showed less seasonal and interperiod variation (Figure 3, top). There was more interperiod variation in daily displacement means relative to net displacement means (Figure 3, top). Correspondingly, measures of variation in the distributions of wild pig movement (e.g., dispersion and scale plots in Figure 3 - middle and bottom) indicated that daily displacement was more variable across periods of the day and season relative to net displacement.

**Figure 3.**
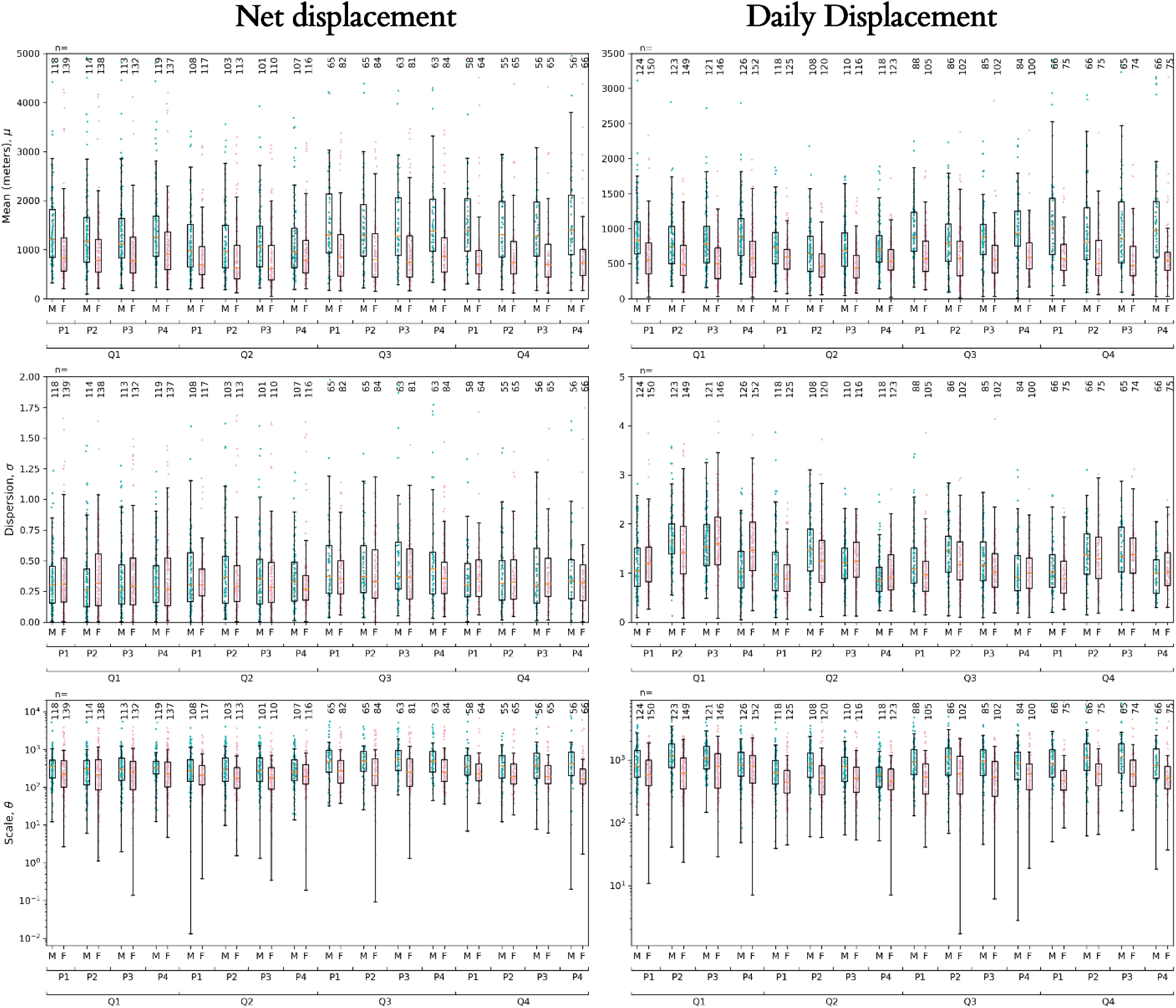
Boxplots of response variables net displacement (left) and daily displacement (right). Each row of two plots summarizes data for different features of the gamma distribution - mean (shape x scale) (top), dispersion (1/scale) (middle), scale (bottom). Y axis units are in meters. X-axes are the categories of period of the day (p1-p4) and season (quarter) of the year (q1-q4). Summaries are also separated for male (blue dots) and female (pink dots) pigs for each period by quarter combination. Number of individual pigs with data for each sex, period, quarter combination are given across the top.

For daily time periods where movement distances were furthest (18:00-6:00), the median for net displacement means ranged from 707 m to 870 m across the four seasons in females, and ranged from 954 m to 1,395 m for males. The mean (**±** 95% confidence interval) of net displacement means ranged from 1,012 **±** 172 m to 1,173 **±** 198 m across the four seasons for females and from 1,210 m to 1,659 m for males. Maxima of net displacement means for females reached 6,616 m for females and 9,417 for males, both occurring during Jan.-Mar. However, mean estimates of net displacement scale (measure of spread in the distribution of individual-level net displacement) were highest during Jan. - Mar. for females and during Oct.-Dec. for males.

The median for daily displacement means across the four seasons in females ranged from 547 m to 585 m and ranged from 703 m to 1,064 m for males. The mean (**±** 95% confidence interval) of daily displacement means ranged from 587 **±** 32 m to 671 **±** 83 m across the four seasons for females and from 758 **±** 48 m to 1,141 **±** 136 m for males. Maxima of daily displacement means for females reached 3,855 m for females and 3,593 m for males, both occurring during Oct.-Dec. Correspondingly, mean estimates of daily displacement scale (measure of spread in the distribution of individual-level daily displacement) were highest during Oct.-Dec. for both females and males.

### Environmental predictors of wild pig net displacement

In general, predictors of net displacement mean and dispersion were more similar across sites than those predicting daily displacement mean and dispersion (Figure 4). The strongest predictors of wild pig net displacement means were variation in the density of secondary roads and mean daylength, followed by variation in primary road density (Figure 4). Net displacement means tended to increase with variation in primary and secondary road density at lower values of the road density but the relationship saturated at higher values (Figure 5). The relationship of net displacement mean to mean daylength was similar to a step function with no effect across most values and increasing strongly at very high values of mean daylength (likely correlated to longer movements in northern latitudes) (Figure 5).

**Figure 4.**
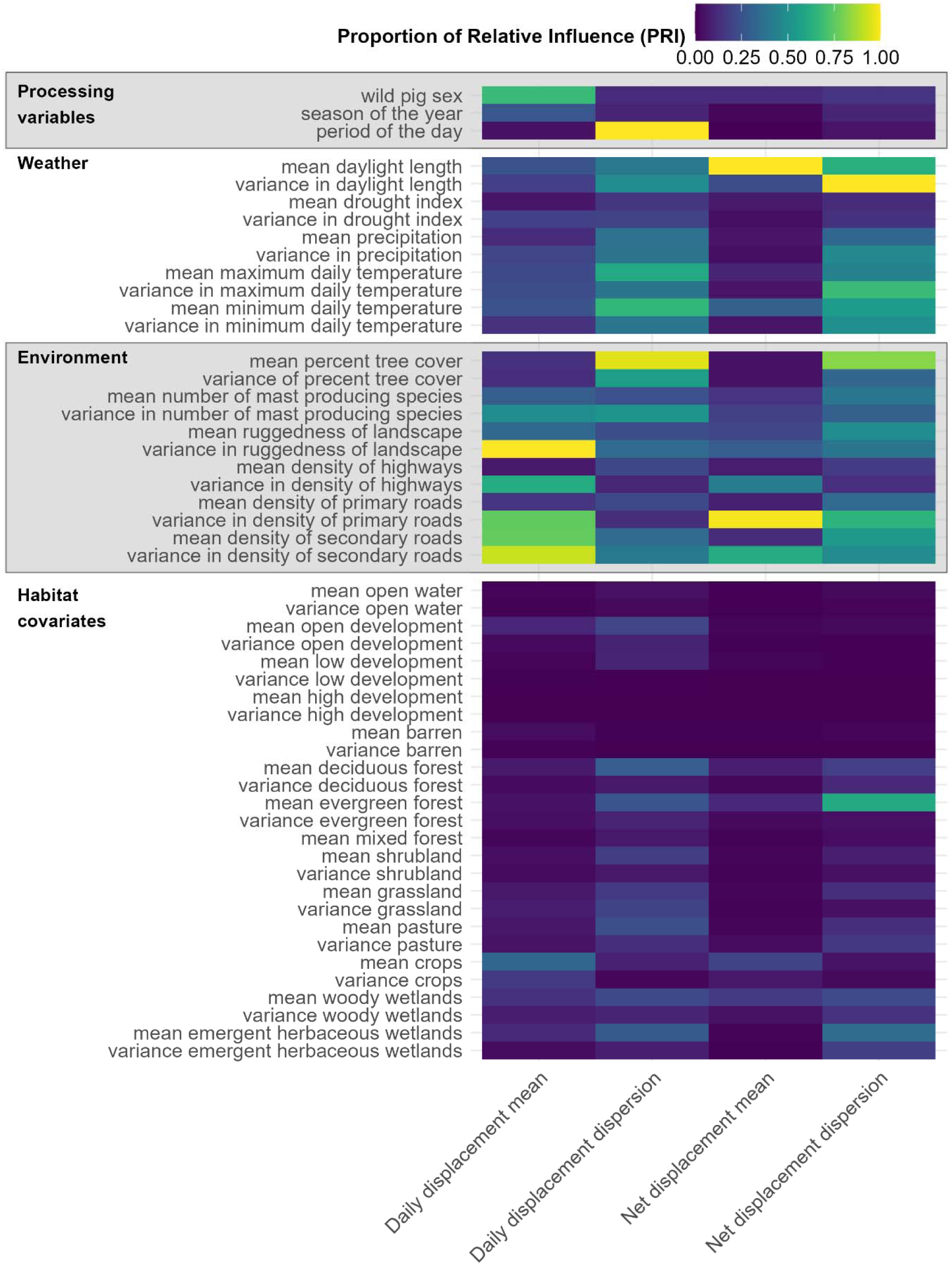
Heat map of proportional relative influence of each predictor on the four response variables. The relative influence of predictors from SGB were scaled to the highest relative influence for each model by dividing all relative influence estimates from a given model by the maximum value for that model. White indicates variables that were dropped because the top model for those responses selected during cross-validation only included variables with the top 90% highest relative influence.

**Figure 5.**
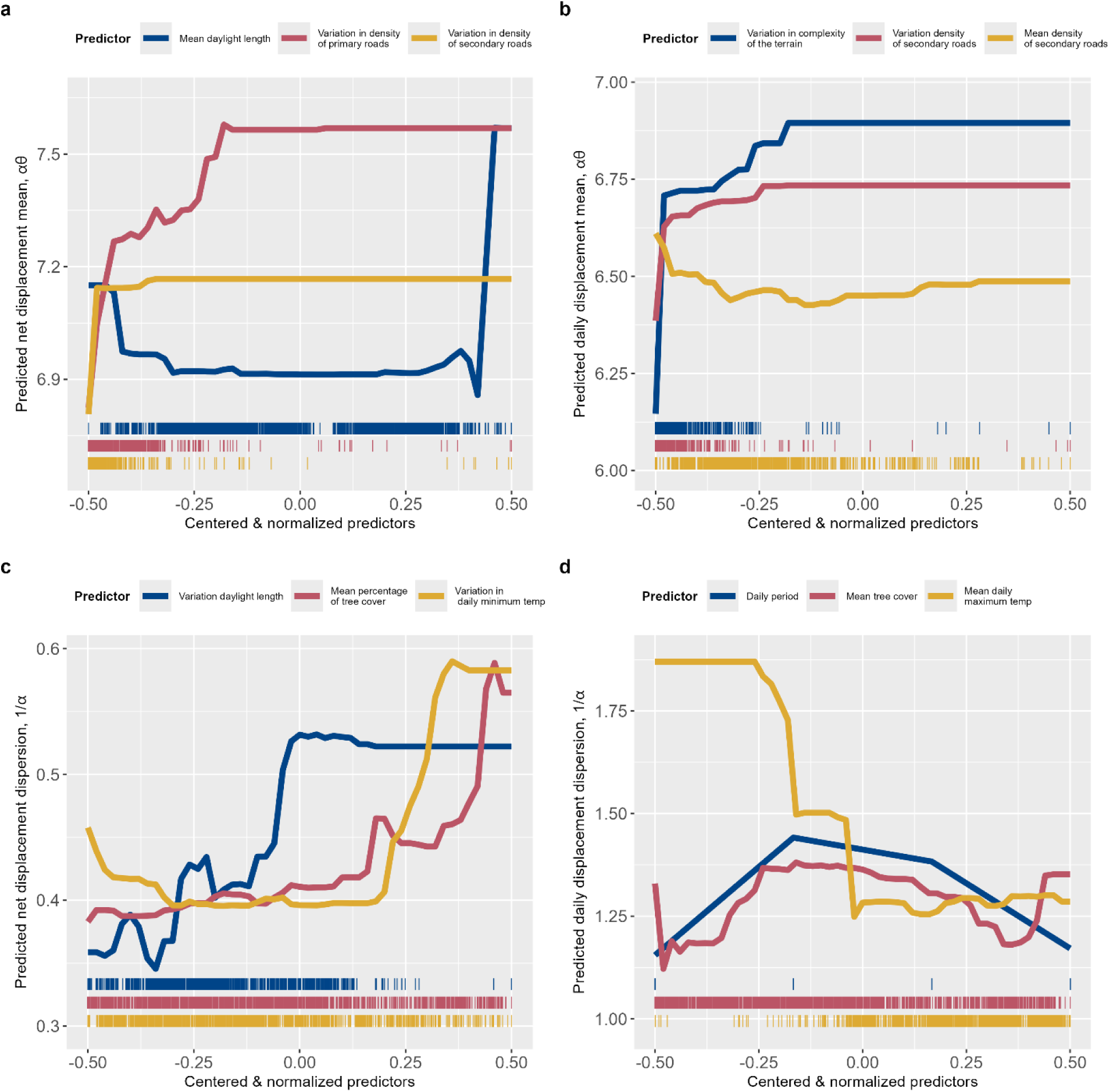
Functional relationships of displacement response variables and some of the strongest predictors. Predictors with the top 3 highest variable importance scores from Figure 4 were plotted for each response variable.

In contrast to net displacement means, there were numerous important predictors of net displacement dispersion, indicating a wide variety of environmental factors driving individual-level variation in extreme values of net displacement distributions (Figure 4). The strongest predictors of the net displacement dispersion included variation in daylength, mean percent of tree cover, and variation in minimum temperature, followed by mean daylength, variation in secondary road density, mean number of mast-producing species, and mean proportion of evergreen species, among others (Figure 4). The relationship of net displacement dispersion to variation in daylength was also similar to a step function with an abrupt shift to higher dispersion values with higher variation in daylength (Figure 5). Net displacement dispersion tended to increase gradually with increasing tree cover proportions, increasing most dramatically at the highest values of tree cover (Figure 5). Net displacement dispersion increased rapidly at higher variation in minimum daily temperature (Figure 5), indicating more extreme values of net displacement at higher extreme values of minimum temperature.

### Environmental predictors of wild pig daily displacement

As with net displacement, a larger number of predictors had high relative importance for daily displacement dispersion relative to daily displacement mean (Figure 4), indicating that a larger number of predictors were important for explaining daily displacement distributions. The strongest predictors of daily displacement means were variation in complexity of the terrain (ruggedness) accessed daily, and the mean and variation of the density of tertiary roads, followed by sex of the individual pig (Figure 4). Daily displacement means increased with terrain complexity to saturation at high values of the predictor (for which there was not a lot of data) and were substantially higher for males relative to females (as found from descriptive summaries in Figure 3). Daily displacement means increased strongly with variation in tertiary road density at lower values (most of the available data), saturating at higher values (Figure 5). But, the relationship of daily displacement means and mean density of tertiary roads was concave up with the highest values of step length being at the lowest and highest values of mean tertiary road density (Figure 5).

Daily displacement dispersion was most strongly predicted by the period of the day (also evident from Figure 3), the mean proportion of tree cover, and the mean maximum daily temperature (Figure 4), followed by the mean minimum daily temperature. Daily displacement dispersion had a complex relationship with mean proportion of tree cover (non-monotonic, generally concave down) but generally decreased with increasing maximum temperature (Figure 5). Daily displacement dispersion tended to be higher during the middle two periods of the day (Figure 5), when daily displacement means were lower on average (Figure 3), indicating there were more extreme net displacement values relative to the average daily displacement during lower movement periods.

### Geographic and seasonal variation in predicted wild pig displacement distances

The model predicted seasonal patterns in the observed data well across the range of wild pigs (Figures 6,7, Figure S2). Variation in predicted net displacement and daily displacement means varied geographically at a fine scale across the wild pig range (Figures 6,7). However, there were also some regional geographic differences. For example, comparing displacement geographically within a season (e.g., Oct-Dec), the average values of predicted net displacement means (all in meters) were higher in more northern states such as Michigan (Mean **±** 1SD: 1,604 **±** 239; Max: 2251), lowest in more southern states such as Florida (Mean **±** 1SD: 1,061 **±** 231; Max: 1,962) and western states such as California (Mean **±** 1SD: 1,065 **±** 174; Max: 1,678), and moderate in southwest states such as Mississippi (Mean **±** 1SD: 1,138 **±** 206; Max: 2,076). Regional averages in daily displacement means with a season (e.g., Oct-Dec) also were apparent, although the trends were different from those of net displacement means. For example, Michigan (Mean **±** 1SD: 845 **±** 127; Max: 1,200) had similar daily displacement means as Florida (Mean **±** 1SD: 756 **±** 145; Max: 1,200), although daily displacement means in another northern state (Wisconsin; where there are not currently any wild pigs) were predicted to be high (Mean **±** 1SD: 1,006 **±** 96; Max: 1,149). Daily displacement means were moderate in central states such as Missouri (Mean **±** 1SD: 809 **±** 127; Max: 1,200) and southeast states such as Mississippi (Mean **±** 1SD: 801 **±** 105; Max: 1,179).

**Figure 6.**
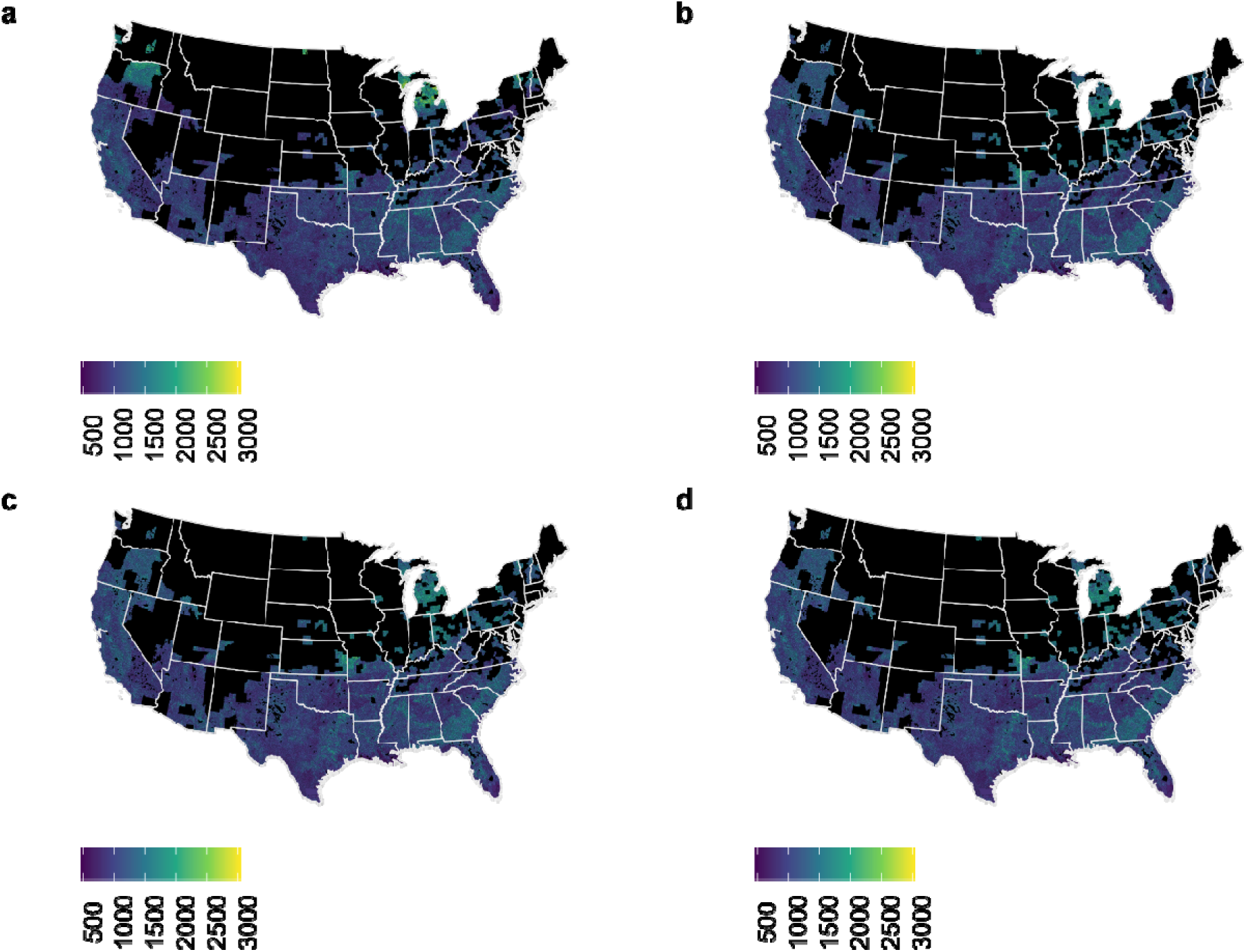
Spatial gradient maps of predicted net displacement means (m) for wild pigs seasonally across the United States. a) January-March; b) April-June; c) July-September; d) October-December. Predictions were made for each watershed across the range where wild pigs have ever been reported according to USDA’s wild pig mapping system.

**Figure 7.**
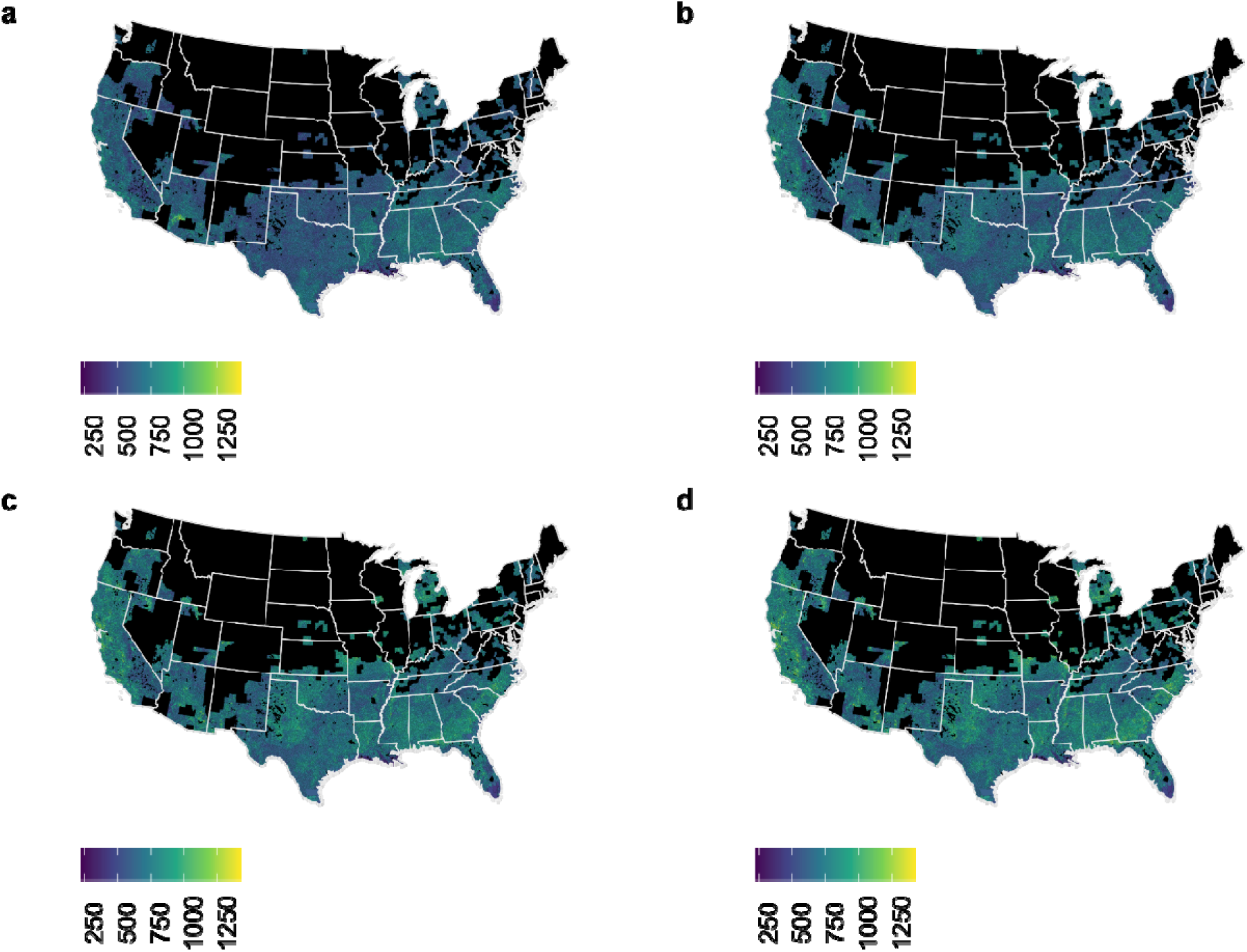
Spatial gradient maps of predicted daily displacement means (m) for wild pigs seasonally across the United States. a) January-March; b) April-June; c) July-September; d) October-December. Predictions were made for each watershed across the range where wild pigs have ever been reported according to USDA’s wild pig mapping system.

### Uncertainty in predicted wild pig displacement distributions

We evaluated the uncertainty for out-of-sample predictions across the range of prediction to identify areas where out-of-sample prediction may be less reliable. Uncertainty in the net displacement means was greater in northern latitudes while uncertainty in net displacement dispersion was higher in Texas and Oklahoma in developed areas along road networks.

Uncertainty in daily displacement means was heterogeneous at small scales, such that many states contained a wide range of uncertainty values. For example, the maximum magnitude of uncertainty was larger for net displacement dispersion, and centered around dense urban areas, while uncertainty around net displacement means was greater at northern latitudes. Regional differences in daily displacement distances (mean and dispersion) were evident, but the drivers were less apparent (Figures S1) and may reflect a combination of environmental variables.

### Habitat selection

Emergent herbaceous wetland was the only habitat type that occurred across most habitat availability groups and consistently had a positive habitat selection coefficient across all habitat availability groups (Figure 8, Table S3–S4). Eight of the twelve groups had access to wetland, consisting of 367 individual pigs (78.9% of all pigs included in the iSSA, Table S5). Barren land also tended to be positive, although it was only present in three habitat availability groups and a small number of individuals. We investigated several of these barren areas using Google maps and discovered they were landfills. Selection for other land cover types depended on habitat availability (Table S4). Habitat selection coefficients for grasslands were generally negative. The direction of selection coefficients for evergreen forests, developed land, pasture, and deciduous forests were strongly dependent on the composition of habitat availability, though not all groups had data from all months (Figure S3). Habitat selection coefficients for woody wetlands were generally neutral for the largest habitat availability group and negative otherwise.

**Figure 8.**
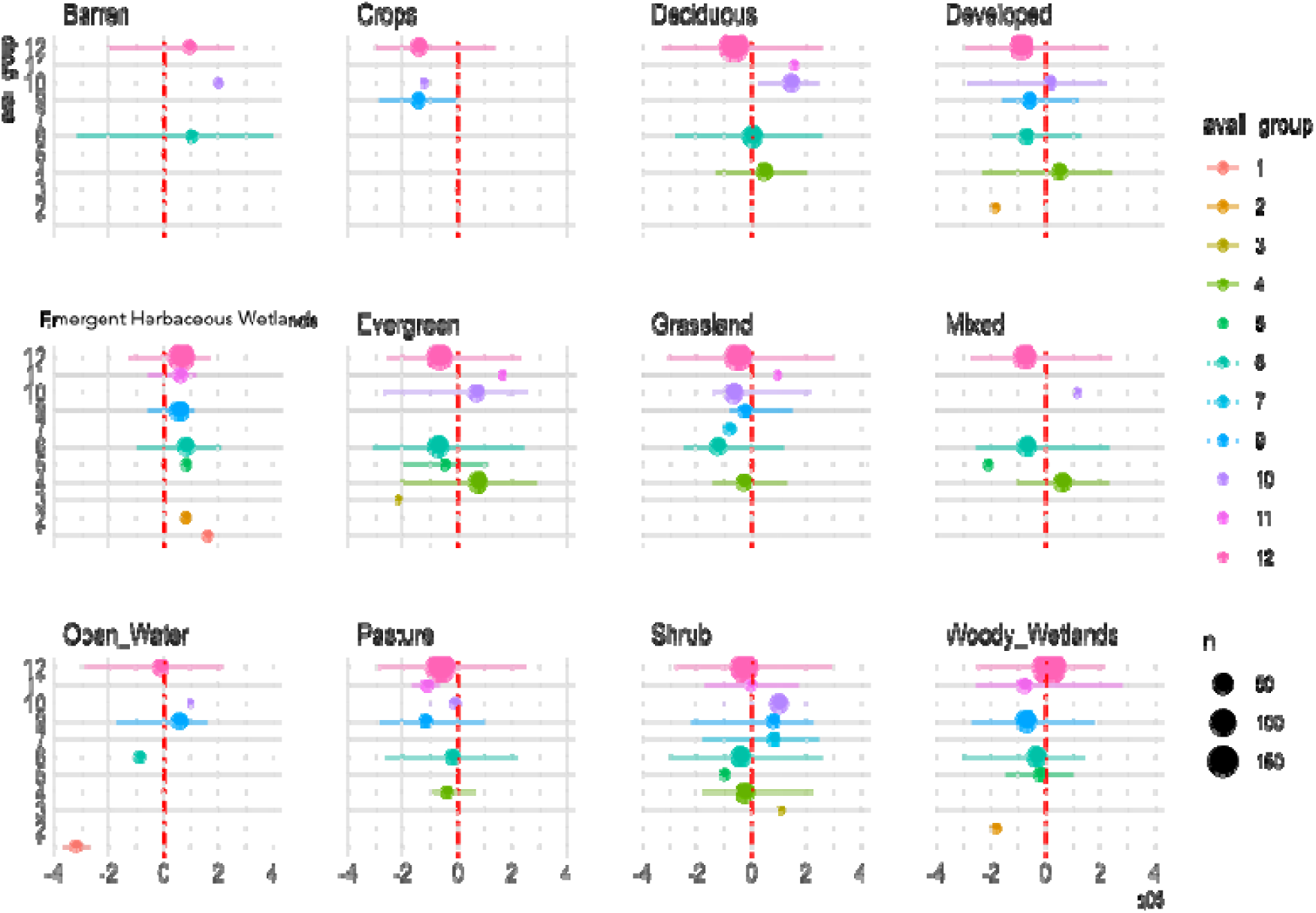
Habitat selection profiles for each group of wild pigs with different habitat availability. Pigs were grouped by habitat availability into 12 groups with the largest group having access to all land types (group 12). Points are sized relative to the number of individual pigs in each group. Points are the iSSA coefficient estimate for each landscape type with standard error

Step length was considered as an interaction term and only seven of the interactions had an effect greater than 1.1 or less than 0.9 preference for every meter moved (Table S6), indicating a very small biological effect on selection. Outside of the group made of an individual pig with access only to open water and emergent herbaceous wetlands, these movement effects were all towards urban (developed) or barren land. Therefore, step length was not included in the final model, as a change of less than 0.1 meter for a meter of movement toward a landscape variable was not considered biologically relevant.

We investigated individuals with regular long movements to verify they were not errors, and the locations visited were confirmed as landfills or a location with food waste, specifically a feed store and a wedding venue with catering. The number of individuals, study sites, total GPS points, and habitat types included in the habitat availability group were positively correlated with the standard errors of the habitat selection coefficient estimates from a leave-one-out analysis (p<0.001 for all four, Figure S4). In general, there was less variation in the habitat availability groups that contained more individuals and included more habitat types (Figure S5–S6). The highest uncertainty in habitat selection coefficients were for shrubs in habitat availability group 7 (2 study sites, 8 individuals, 6 habitat types) and crops and evergreen forests in habitat availability 11 (6 study sites, 13 individuals, 10 habitat types), demonstrating limitations from sample size, especially in the less frequently seen landscapes type subsets.

## Discussion

We characterized heterogeneity in wild pig movement behavior across the species’ range in North America and examined landscape correlates with these movement properties with the aim of enabling range-wide (i.e., out of sample) predictions of wild pig movement metrics. Our work focused on key movement metrics at a coarser temporal scale (daily) than is typical for high-resolution GPS data to provide inferences that are relevant to downstream applications such as disease management decision-making and population control. Providing estimates of movement metrics in unsampled areas allows for predicting population expansion and disease spread rates anywhere in the country. Our analysis revealed new insights about how natural wild pig movement behavior varies geographically and seasonally across their range in the contiguous U.S. through relationships with local environmental variables.

### Observed trends in movement distances

Several overall patterns for movement were observed across all datasets. Male displacement was further and more variable in annual quarters three and four relative to the first half of the year, and greater than that of females across the year. Larger home ranges and further displacement by males is consistently reported in wild pigs across multiple studies within the continental United States and in other regions outside the range of the current study (continental United States) [40][67,68]. Increases in net and daily displacement for males during annual quarters three and four could be associated with increased movements of males to find reproductively receptive females. Previous studies in North America have reported an increase in farrowing activity from November to May which would correspond to increases in conception during the fall and early winter [69]. Additionally, other studies have proposed that larger winter and spring home ranges are associated with lower forage availability requiring greater movement [70]. Others have found the opposite and suggested that increased food availability may actually increase movement, such as mast availability in fall [40,71], or found no difference seasonally [72]. Females had less seasonal and interperiod variation in net and daily displacement relative to males. Overall, daily displacement shows more variation seasonally and across periods of the day than net displacement for both males and females. Maxima of net displacement means at the daily scale for females reached 6.6 km and 9.4 km for males, both occurring during the first quarter. In comparison, maxima of daily displacement means for females reached 3.9 km for females and 3.6 km for males, both occurring during the fourth quarter. Extreme movements can drive disease transmission and range expansion of invasive species [73]. Thus, these large movements are of importance to better understand and predict disease spread for management purposes. While our work highlights the greater distances moved by males, it also highlights the more consistent mobility of females, information important to structure monitoring efforts or interventions.

### Effects of environmental predictors

Predictors of net displacement means and dispersion were more similar across sites than those predicting daily displacement means and dispersion, suggesting that individual-level factors or localized dynamics affected daily displacement more strongly than net displacement. As such, net displacement may be predicted more strongly by factors occurring at broader spatial scales, rather than local or inter period conditions. One of the strongest, positive influences on net displacement means was variation in road density. Roads can facilitate movement for feral swine by allowing for easier travel across terrain or, conversely, inhibit movement by acting as a barrier to movement when experiencing high traffic volumes [40][72][74]. Thus, when availability of roads is more variable there is likely increased variation in the juxtaposition of wild lands and roads. Wild pigs may exhibit longer net displacement distances as they access roads that facilitate travel.

Mean daylength was an important predictor of net displacement but only influenced net displacement at very high values of mean daylength, which is likely correlated to longer movements in northern latitudes during the summer. Studies at very northern latitudes (e.g., Sweden) have found a similar increase in movement during longer summer days [74]. There were several strong predictors of net displacement dispersion. Our finding that variation in minimum temperature influenced net displacement dispersion is consistent with previously reported relationships of movement and abiotic conditions [75]. Also, our finding that increased net and daily displacement dispersion were related to percent tree cover aligns with the fact that mast resources change seasonally, and shifts in masting resource quality can vary spatially and temporally [76,77]. Because higher variability in movement distance could drive disease spread more rapidly over longer distances [24,73,78], this may mean that areas where individuals are most reliant on mast-producing resources and times of the year where these resources are highly variable could be important considerations for emergency disease management as individuals may shift movement in these conditions. Other studies have also suggested greater movement in response to mast availability [71].

In comparison, the strongest predictors of daily displacement means were variation in terrain ruggedness, mean and variation of the density of tertiary roads, followed by sex of the individual pig. Terrain ruggedness increases step length, likely through increasing the distance animals have to move between preferred lower gradient slopes. A study in the Great Smoky Mountains found pigs preferred flat areas year-round and shifted to lower elevation in the winter [79]. Daily displacement dispersion had similar types of predictors as net displacement dispersion: period of the day, tree cover, and temperature. In many areas, feral swine are diurnal and in some regions and times of the year can be nocturnal [65,80,81]. This daily pattern of activity has been reported to be affected by a variety of abiotic and anthropogenic factors. Swine may shift to more nocturnal activity in response to anthropogenic threats such as daylight hunting or management related to crop protection [71,82][83]. Temperature has also been associated with greater diurnal activity, with a shift to movement during cooler crepuscular and nocturnal temperatures in certain sites [84]. Wild pigs decreased activity with increased maximum temperature in multiple studies [40][81]. However, not all studies found a decrease in activity with increased temperature in the summer but just with increasing temperature in the early growing season [46]. The effect of increased temperature on decreasing wild pig movement likely interacts with available thermal habitats such as woody wetlands [82,85] and is more prominent at higher extremes for maximum temperature.

### Geographic differences in predictions

Our spatial predictions identified ‘hot spots’ of longer daily displacement means, with high variability at finer spatial scales. Under this model framework, these differences in spatial variation reflect the spatial variation in environmental covariates with the highest percent relative influence. In contrast, we predicted coarse spatial variation in net displacement means, with a clear positive association with higher latitudes. Some of the strongest environmental drivers of daily displacement were variation in terrain complexity, and mean and variance of tertiary road density. The greatest shifts in net displacement means were related to primary and secondary road density, as well as mean daylight length. Unlike the terrain complexity and road density variables, which are more variable at finer spatial scales, mean daylight length varies across broad latitudinal ranges. Thus, predictions that rely on this variable are likely to vary across broad ranges as well. While this study has encompassed the largest diversity of locations in the

U.S. to date, some similar patterns have emerged that were previously reported in studies using only data from the southern U.S. Our finding that pigs in subtropical parts of the U.S. (i.e. Florida) have the shortest movement distances is similar to [40]. This may be due to higher productivity in subtropical habitats. However, we did not find greater movements in Texas relative to Florida reported by [40]. This is likely due to site specific heterogeneity across Texas. Our data included additional sites from Texas that were not included in [40] resulting in mean movement distances for Texas that are similar to those in Florida. We saw further movements in the more northern states and moderate movement in the southeast.

### Habitat selection

Our iSSA results indicated that habitat selection profiles depend on ecological context. We identified general and divergent selection for specific habitat across differing ecological contexts by grouping individuals by available land cover types. Emergent herbaceous wetland was the only habitat type that occurred across most habitat availability groups and consistently had a positive habitat selection coefficient across all habitat availability groups. This is important as feral swine are known to damage fragile wetland ecosystems with their rooting behavior [60,66]. Other studies have suggested that wetlands may also serve for thermoregulation alongside being a food source [68].

In comparison, habitat selection coefficients for grasslands were generally negative. Habitat selection coefficients for woody wetlands were generally neutral for the largest habitat availability group and negative otherwise. This nuance would have been lost if the full data were analyzed all together across the same set of habitat availability. Our analysis likely underestimated preference for crop land because attraction to crop land is specific to crop type and sometimes during short time windows during the year (i.e., during planting or harvesting) [63]. Few individual pigs had access to the barren land type category but those that did showed a preference for this land type likely because the barren land cover in our study tended to be landfills. Previous work has demonstrated that landfills are attractants for wild pigs [86–88]. However, a previous study using much of the same data as this study found that pigs further away from crops did not move towards crops suggesting pigs residing further away from crops, may not travel to the crops routinely [40]. Furthermore, in baboons, another social crop-raiding species, troops are found to spend more time adjacent to fields as compared to actively raiding [89] so the temporal scale used for this study may affect observations of crop raiding activity.

### Caveats and generalizability

Characterizing uncertainty is important in forecasting studies to provide guidance for the level of confidence that can be placed in a given prediction across the range of predictions [90,91]. We quantified spatial uncertainty in movement predictions based on ensemble model disagreement [55,57,58]. High uncertainty aligned with regions containing covariate combinations poorly represented in our training data. For example, we found high uncertainty of net displacement predictions in regions with high levels of developed land and primary road density and in northern latitudes, because most of our relocation data did not overlap these habitat features. Our dataset did not contain individuals in urban areas, and the only data we evaluated from northern latitudes was from northern Michigan. Alternative approaches like Multivariate Environmental Similarity Surfaces (MESS; [92]) could complement our regional uncertainty characterization by directly linking prediction uncertainty to specific covariates.

Ultimately, out-of-sample spatial prediction accuracy depends on both sample size (relative to the scope of prediction) and covariate coverage of the training data [93]. Sensitivity analyses examining how sample size affects model accuracy have been a common approach to assessing this relationship in species distribution models [94,95]. Methods that jointly assess sample size and predictor coverage have been developed in clinical settings [96]. For species distribution models, sample size matters more than spatial sampling bias [93]. Evaluating optimal sample size and covariate coverage needed for accurate movement predictions could improve this framework.

Our analysis captured the central tendency and variation features of the distribution of wild pigs well, thus providing reasonable summaries of the most commonly observed patterns of net displacement mean and dispersion. However, there were infrequent extreme values that were not well captured by our approach. This may be partly due to our choice of a gamma distribution for summarizing net and daily displacement. While gamma distributions have some skew they are unimodal and described by only two parameters, introducing some limitations for capturing extreme values. Thus for species where extreme values are more frequent (e.g., migratory species) a more complex unimodal distribution, mixture distribution, or non-parametric distribution may be preferable (e.g., [97,98]). Also, because longer-range movements may occur on a different timescale, it may be also important to predict their timing (e.g., [99]). Thus, another approach could be to separate ‘usual’ movement distances from longer-range movements and develop separate models that predict their values. Ultimately, choosing the time scale and functional form(s) for modeling the distributions of displacements should depend on the species movement ecology and downstream application of interest. For our application of approximating wild pig movement distances as input parameters to emergency disease management [24], our analysis demonstrates that a gamma distribution demonstrates a reasonable, parsimonious summary of the relevant variation in natural wild pig displacement distances. However, long-distance translocation of wild pigs [34] is an additional consideration that may affect disease spread, but predicting the distance or direction of translocations [100], or wild pig movement behavior following translocation [101,102], from environmental drivers is challenging.

There were some further limitations in our iSSA modeling. Due to the variance in GPS-collar fix rates, we had to resample all datasets to the lowest frequency rate, which was every six hours. This meant the level of detail was equal over each study, but some information was lost. This is a consistent and known issue with upscaling data at the pre-model stage [103] and is a common limitation when modeling geolocation data. Lastly, the variability in effect size was evident for habitat availability groups containing fewer samples. Potentially, as these land cover combinations occurred more rarely in our data set, they may be less relevant on a national scale. In addition, seasonal effects were not examined for the iSSA model as not all groupings had data from the entire year. Seasonal differences may be especially relevant for crop use, as seasonality affects the value of some crops that can serve as a source of protective cover as well as food [63,104]. This may further explain why crops were frequently selected against in the iSSA model. The negative selection of crops might change if seasonality or crop types were incorporated [36].

Our framework predicts aggregate movement characteristics across spatial extents where local movement studies may not exist, making it suitable for landscape scale management requiring broad movement characterization such as barrier placement or buffer zone planning (e.g., disease control containment zones; [105]). This framework is less suitable for management applications requiring behaviorally specific movement characteristics (i.e., focusing on juvenile dispersal). For example, landscape interventions may require different considerations for migratory versus home range movements [106]. Future versions of this framework could improve application for these systems by integrating behavioral state classification into the modeling pipeline by first classifying trajectories into behavioral states (e.g., [107]), then modeling them separately or as covariates. This would enable comparison of movement drivers across behavioral states and better support ecological research or impact assessments requiring phenologically-explicit movement understanding.

## Conclusions

We predicted net and daily displacement and habitat selection of individual wild pigs across wide-ranging ecological contexts, quantifying variation in movement behavior across the species’ range. Our approach provided novel insights into wild pig movement behavior and provided quantitative assessment of the effects of environmental factors on wild pig movement. Our predictions provide parameters for informing ecological models of disease spread or population invasion for informing downstream management applications. Our approach can be adapted to finer temporal scales (e.g., hourly steps) or coarser scales (e.g., weekly or monthly movement metrics) depending on need for the ecological question or management application.

## Declarations

### Ethics approval and consent to participate

Not applicable

## Consent for publication

Not applicable

## Data availability statement

Data and code used to generate these results will be provided upon publication. All geolocation data provided has been anonymized by transforming the coordinates. For reviewers and editors, code and data to run model are uploaded to figshare at: figshare.com/s/14c31c3d84eb793f2234?file=54273221

## Acknowledgements

The authors thank multiple collaborators who shared wild pig geolocation data including Susan M. Cooper, Tyler Campbell, Lindsay Holmstrom, Steve Hartley, and Michael D. White. We also acknowledge the contributions of Jonathan Potts, Luca Börger, Bronson K. Strickland, and Garrett M. Street whose publicly available wild pig geolocation data were used in this work. The authors also thank Josh Hewitt for feedback on the modeling approach. The findings and conclusions in this publication are those of the authors and should not be construed to represent any official US Government determination or policy. Mention of commercial products or companies does not represent an endorsement by the US government.

## Supplementary Information

**Table S1.**
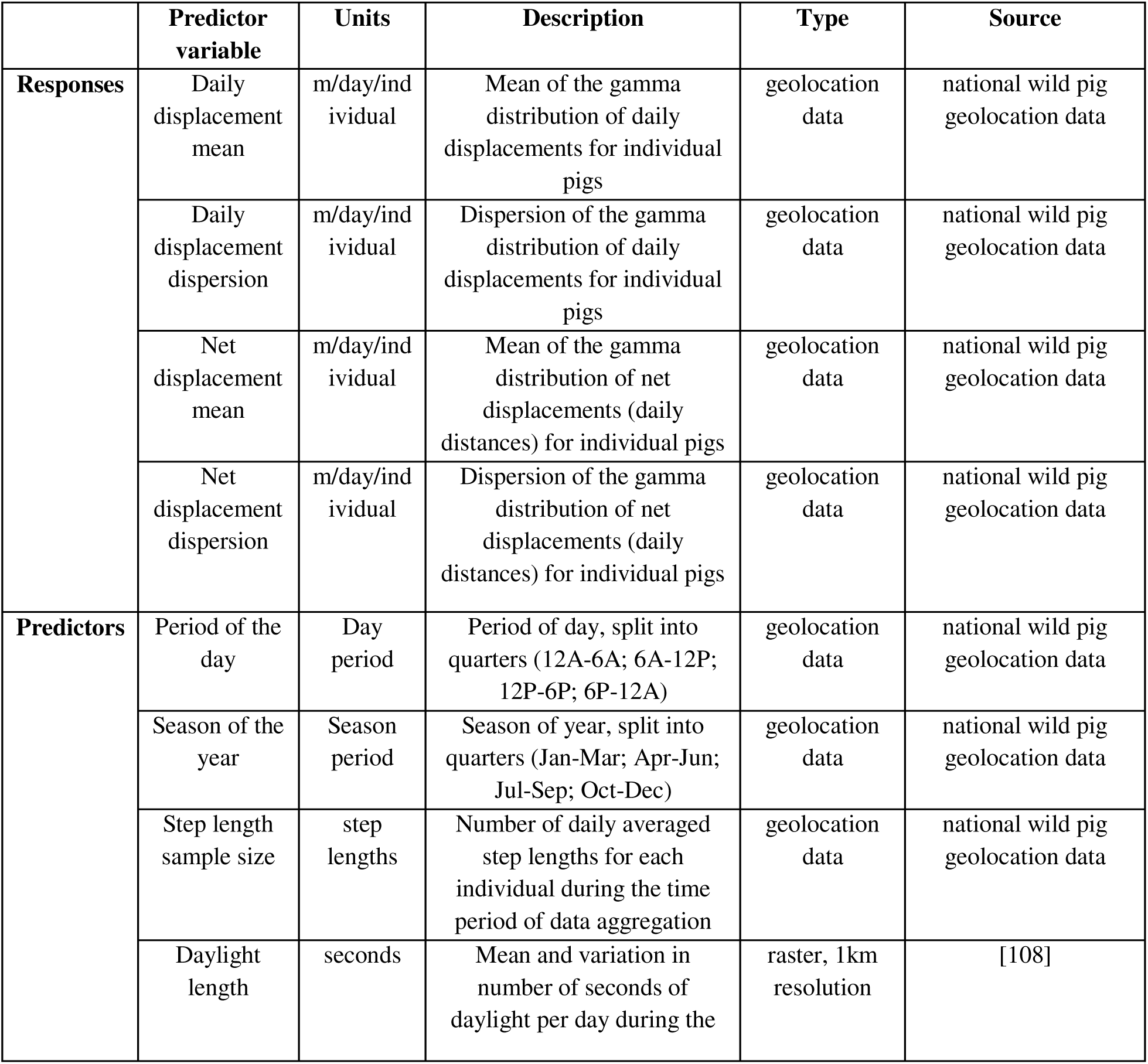
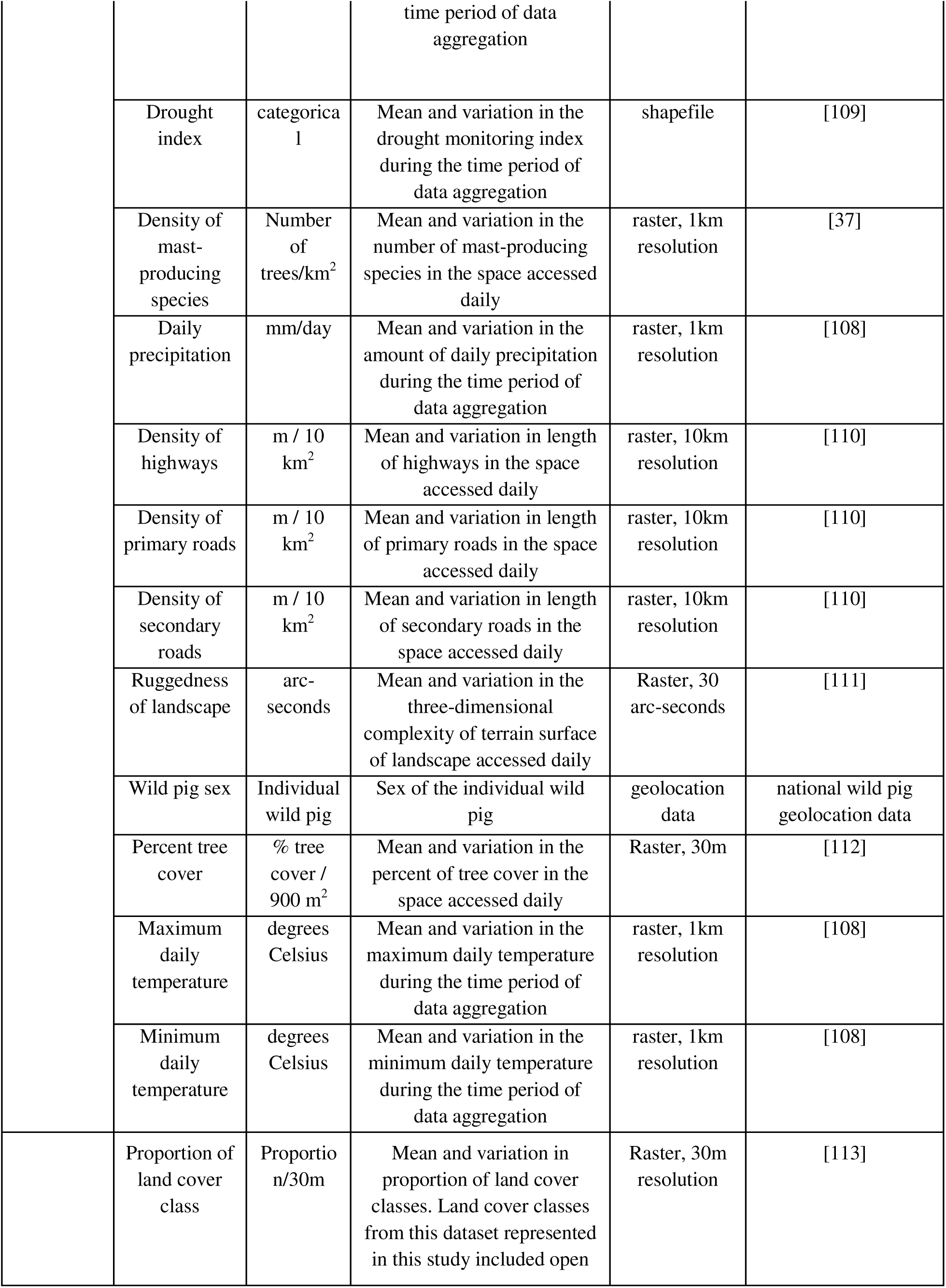
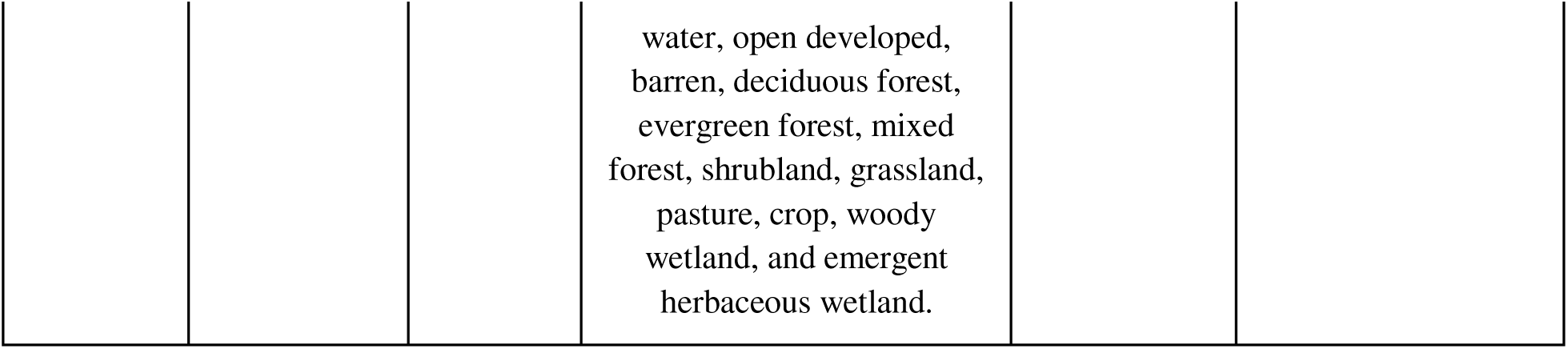
Description of response and predictor variables for movement distance metrics.

|  | Predictor variable | Units | Description | Type | Source |
| --- | --- | --- | --- | --- | --- |
| **Responses** | Daily displacement mean | m/day/individual | Mean of the gamma distribution of daily displacements for individual pigs | geolocation data | national wild pig geolocation data |
|  | Daily displacement dispersion | m/day/individual | Dispersion of the gamma distribution of daily displacements for individual pigs | geolocation data | national wild pig geolocation data |
|  | Net displacement mean | m/day/individual | Mean of the gamma distribution of net displacements (daily distances) for individual pigs | geolocation data | national wild pig geolocation data |
|  | Net displacement dispersion | m/day/individual | Dispersion of the gamma distribution of net displacements (daily distances) for individual pigs | geolocation data | national wild pig geolocation data |
| **Predictors** | Period of the day | Day period | Period of day, split into quarters (12A-6A; 6A-12P; 12P-6P; 6P-12A) | geolocation data | national wild pig geolocation data |
|  | Season of the year | Season period | Season of year, split into quarters (Jan-Mar; Apr-Jun; Jul-Sep; Oct-Dec) | geolocation data | national wild pig geolocation data |
|  | Step length sample size | step lengths | Number of daily averaged step lengths for each individual during the time period of data aggregation | geolocation data | national wild pig geolocation data |
|  | Daylight length | seconds | Mean and variation in number of seconds of daylight per day during the | raster, 1km resolution | [108] |

|  |  |  |  |  |
| --- | --- | --- | --- | --- |
|  |  | time period of data aggregation |  |  |
| Drought index | categorical | Mean and variation in the drought monitoring index during the time period of data aggregation | shapefile | [109] |
| Density of mast-producing species | Number of trees/km<sup>2</sup> | Mean and variation in the number of mast-producing species in the space accessed daily | raster, 1km resolution | [37] |
| Daily precipitation | mm/day | Mean and variation in the amount of daily precipitation during the time period of data aggregation | raster, 1km resolution | [108] |
| Density of highways | m / 10 km<sup>2</sup> | Mean and variation in length of highways in the space accessed daily | raster, 10km resolution | [110] |
| Density of primary roads | m / 10 km<sup>2</sup> | Mean and variation in length of primary roads in the space accessed daily | raster, 10km resolution | [110] |
| Density of secondary roads | m / 10 km<sup>2</sup> | Mean and variation in length of secondary roads in the space accessed daily | raster, 10km resolution | [110] |
| Ruggedness of landscape | arc-seconds | Mean and variation in the three-dimensional complexity of terrain surface of landscape accessed daily | Raster, 30 arc-seconds | [111] |
| Wild pig sex | Individual wild pig | Sex of the individual wild pig | geolocation data | national wild pig geolocation data |
| Percent tree cover | % tree cover / 900 m<sup>2</sup> | Mean and variation in the percent of tree cover in the space accessed daily | Raster, 30m | [112] |
| Maximum daily temperature | degrees Celsius | Mean and variation in the maximum daily temperature during the time period of data aggregation | raster, 1km resolution | [108] |
| Minimum daily temperature | degrees Celsius | Mean and variation in the minimum daily temperature during the time period of data aggregation | raster, 1km resolution | [108] |
| Proportion of land cover class | Proportion/30m | Mean and variation in proportion of land cover classes. Land cover classes from this dataset represented in this study included open | Raster, 30m resolution | [113] |

|  |  |  |  |
| --- | --- | --- | --- |
|  |  |  | water, open developed,<br>barren, deciduous forest,<br>evergreen forest, mixed<br>forest, shrubland, grassland,<br>pasture, crop, woody<br>wetland, and emergent<br>herbaceous wetland. |

**Table S2.**
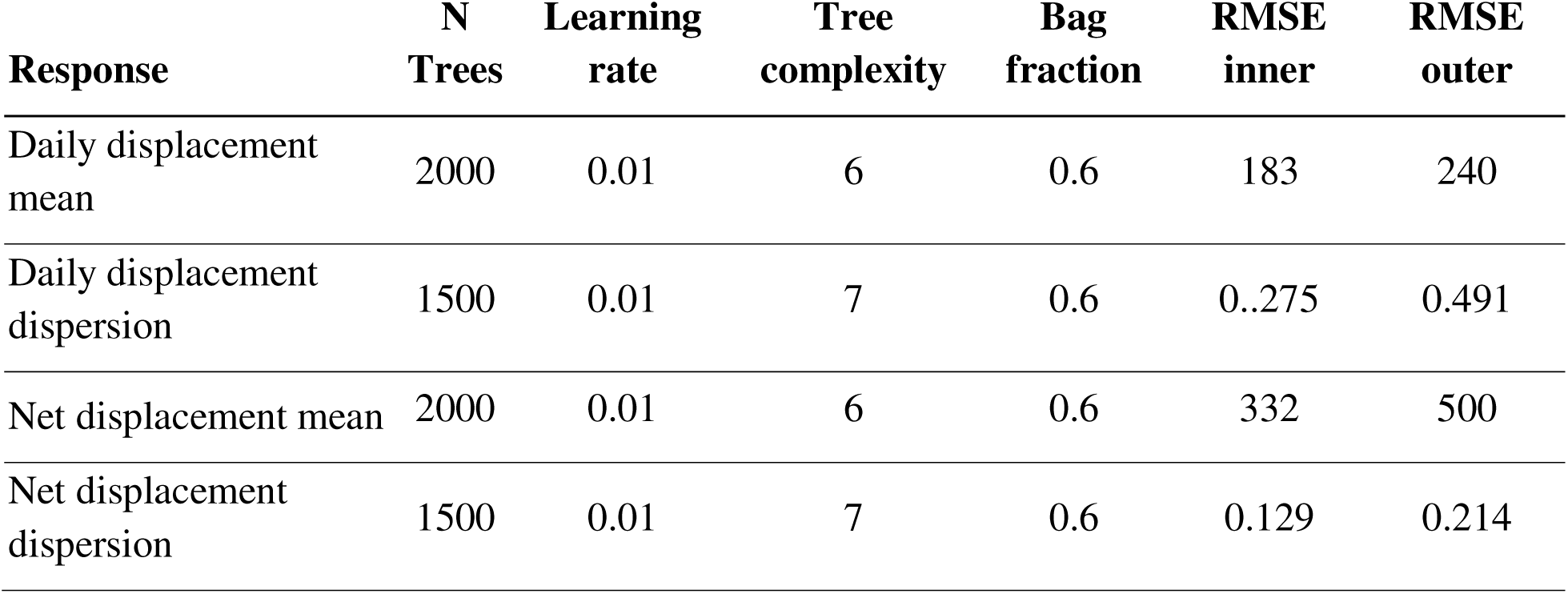
Optimal hyperparameters for fitting response variables. N trees- number of trees.

| **Response** | **N**<br>**Trees** | **Learning**<br>**rate** | **Tree**<br>**complexity** | **Bag**<br>**fraction** | **RMSE**<br>**inner** | **RMSE**<br>**outer** |
| --- | --- | --- | --- | --- | --- | --- |
| Daily displacement<br>mean | 2000 | 0.01 | 6 | 0.6 | 183 | 240 |
| Daily displacement<br>dispersion | 1500 | 0.01 | 7 | 0.6 | 0.275 | 0.491 |
| Net displacement mean | 2000 | 0.01 | 6 | 0.6 | 332 | 500 |
| Net displacement<br>dispersion | 1500 | 0.01 | 7 | 0.6 | 0.129 | 0.214 |

**Table S3.** Mean estimates from individuals in which the estimate was significant from step selection model for different land types, subset according to access to wetlands, urban areas, forests, and cropland (n=464).

| **Subset**<br>**Number**<br>**(NLCD**<br>**number)** | **1** | **2** | **3** | **4** | **5** | **6** | **7** | **8** | **9** | **10** | **11** | **12** |
| --- | --- | --- | --- | --- | --- | --- | --- | --- | --- | --- | --- | --- |
| open water<br>(11) | -3.22 |  |  |  |  | -0.88 |  |  | 0.58 | 0.99 |  | -0.10 |
| developed<br>(21) |  | -1.83 |  | 0.50 |  | -0.70 |  |  | -0.58 | 0.16 |  | -0.89 |
| barren (31) |  |  |  |  |  | 1.03 |  |  |  | 2.01 |  | 0.95 |
| deciduous<br>forest (41) |  |  |  | 0.47 |  | 0.84 |  |  |  | 1.44 | 1.54 | -0.64 |
| evergreen<br>forest (42) |  |  | -2.16 | 0.76 | -0.45 | -0.70 |  |  |  | 0.68 | 1.62 | -0.66 |
| mixed forest<br>(43) |  |  |  | 0.62 | -2.10 | -0.68 |  |  |  | 1.17 |  | -0.75 |
| shrubland<br>(52) |  |  | 1.08 | -0.25 | -0.99 | -0.41 | 0.81 |  | 0.79 | 1.00 | -0.03 | -0.29 |
| grassland (71) |  |  |  | -0.27 |  | -1.24 | -0.82 |  | -0.24 | -0.64 | 0.95 | -0.50 |
| pasture (81) |  |  |  | -0.38 |  | -0.19 |  |  | -1.18 | -0.10 | -1.10 | -0.62 |
| crops (82) |  |  |  |  |  |  |  |  | -1.43 | -1.21 |  | -1.41 |
| woody wetlands (90) |  | -1.80 |  |  | 0.19 | -0.35 |  |  | -0.71 |  | -0.79 | 0.14 |
| emergent herbaceous wetland (95) | 1.61 | 0.82 |  |  | 0.83 | 0.83 |  |  | 0.58 |  | 0.61 | 0.68 |

**Table S4.** Median estimates from ALL individuals from step selection model for different land types, subset according to access to wetlands, urban areas, forests, and cropland (n=464).

| Subset Number | 1 | 2 | 3 | 4 | 5 | 6 | 7 | 8 | 9 | 10 | 11 | 12 |
| --- | --- | --- | --- | --- | --- | --- | --- | --- | --- | --- | --- | --- |
| open water (11) | -3.22 | -0.73 |  |  | -0.37 | -0.22 | 0.18 | -0.21 | 0.38 | 0.34 | -0.14 | -0.15 |
| developed (21) |  | -1.31 |  | 0.31 |  | -0.52 | 0.20 |  | -0.08 | 0.39 |  | -0.42 |
| barren (31) |  |  |  |  |  | 0.40 |  |  |  | 1.49 | 0.01 | -0.14 |
| deciduous forest (41) |  |  | -0.12 | 0.32 |  | -0.24 |  |  |  | 0.49 | 0.24 | -0.41 |
| evergreen forest (42) |  |  | -0.56 | 0.10 | -0.06 | -0.59 |  |  |  | 0.53 | 0.33 | -0.18 |
| mixed forest (43) |  |  | 0.32 | 0.02 | -0.61 | -0.40 |  |  |  | -0.06 |  | -0.15 |
| shrubland (52) |  |  | 0.13 | -0.09 | -0.02 | NA | -0.18 | 4.47 | 0.75 | 0.66 | -0.73 | -0.23 |
| grassland (71) |  |  | -0.06 | -0.01 |  | NA | -0.85 |  | 0.20 | -0.33 | 0.40 | -0.29 |
| pasture (81) |  |  |  | -0.18 | 0.00 | -0.64 | 13.33 | 0.15 | -0.26 | 0.12 | -0.03 | -0.22 |
| crops (82) |  |  |  |  |  |  | -0.12 |  | -0.49 | -0.28 | -0.67 | -0.37 |
| woody wetlands (90) |  | 1.54 |  |  | 0.09 | -0.35 |  | -0.82 | -0.31 |  | -0.67 | 0.05 |
| emergent herbaceous wetland (95) | 1.61 | -0.64 |  |  | 0.05 | 0.80 |  | 0.02 | 0.21 |  | -0.12 | 0.25 |

**Table S5.** Presence of Key Land Types for Each Habitat Availability Subset (n=464)

| **Subset Number** | **1** | **2** | **3** | **4** | **5** | **6** | **7** | **8** | **9** | **10** | **11** | **12** |
| --- | --- | --- | --- | --- | --- | --- | --- | --- | --- | --- | --- | --- |
| Crop available |  |  |  |  |  |  | 1 | 1 | 1 | 1 | 1 | 1 |
| Urban available |  | 1 |  | 1 |  | 1 | 1 |  | 1 | 1 |  | 1 |
| Wetland available | 1 | 1 |  |  | 1 | 1 |  | 1 | 1 |  | 1 | 1 |
| Forest available |  |  | 1 | 1 | 1 | 1 |  |  |  | 1 | 1 | 1 |
| Pigs (n) | 1 | 4 | 2 | 52 | 8 | 71 | 8 | 2 | 49 | 35 | 13 | 219 |
| Number of study sites per group<br>(n=23) | 1 | 1 | 2 | 6 | 4 | 12 | 2 | 2 | 8 | 5 | 6 | 23 |

**Table S6.** Comparison of step selection model for different land types with and without step length (SL) as an interaction term, subset according to access to wetlands, urban areas, forests, and cropland.

| Group Number | 1 | 2 | 3 | 4 | 5 | 6 | 7 | 8 | 9 | 10 | 11 | 12 |
| --- | --- | --- | --- | --- | --- | --- | --- | --- | --- | --- | --- | --- |
| Change in AIC | -11.3 | -0.41 | 3.81 | -70.7 | -17.55 | -36.1 | 6.18 | 5.61 | -300.04 | -36.03 | -11.18 | -562.41 |
| Number of significant interactions with SL | 1 | 1 | 0 | 4 | 5 | 7 | 0 | 0 | 4 | 1 | 2 | 6 |
| Largest significant effect size (p) | 5.25<br>(<0.001) | 1.28<br>(0.04) |  | 1.00<br>(<0.001) | 0.99<br>(<0.001) | 1.92<br>(<0.001) |  |  | 1.23<br>(<0.001) | 0.25<br>(<0.001) | 1.80<br>(<0.001) | 1.59<br>(<0.001) |
| Land type with significant estimates of >1.1 or <0.9* | 95 | 21 |  |  |  | 21 |  |  | 21 | 21 | 31 | 21 |

\*NLCD Key: 95 = Emergent Herbaceous Wetlands , 21= Developed (all intensities combined), 31=Barren

**Figure S1.**
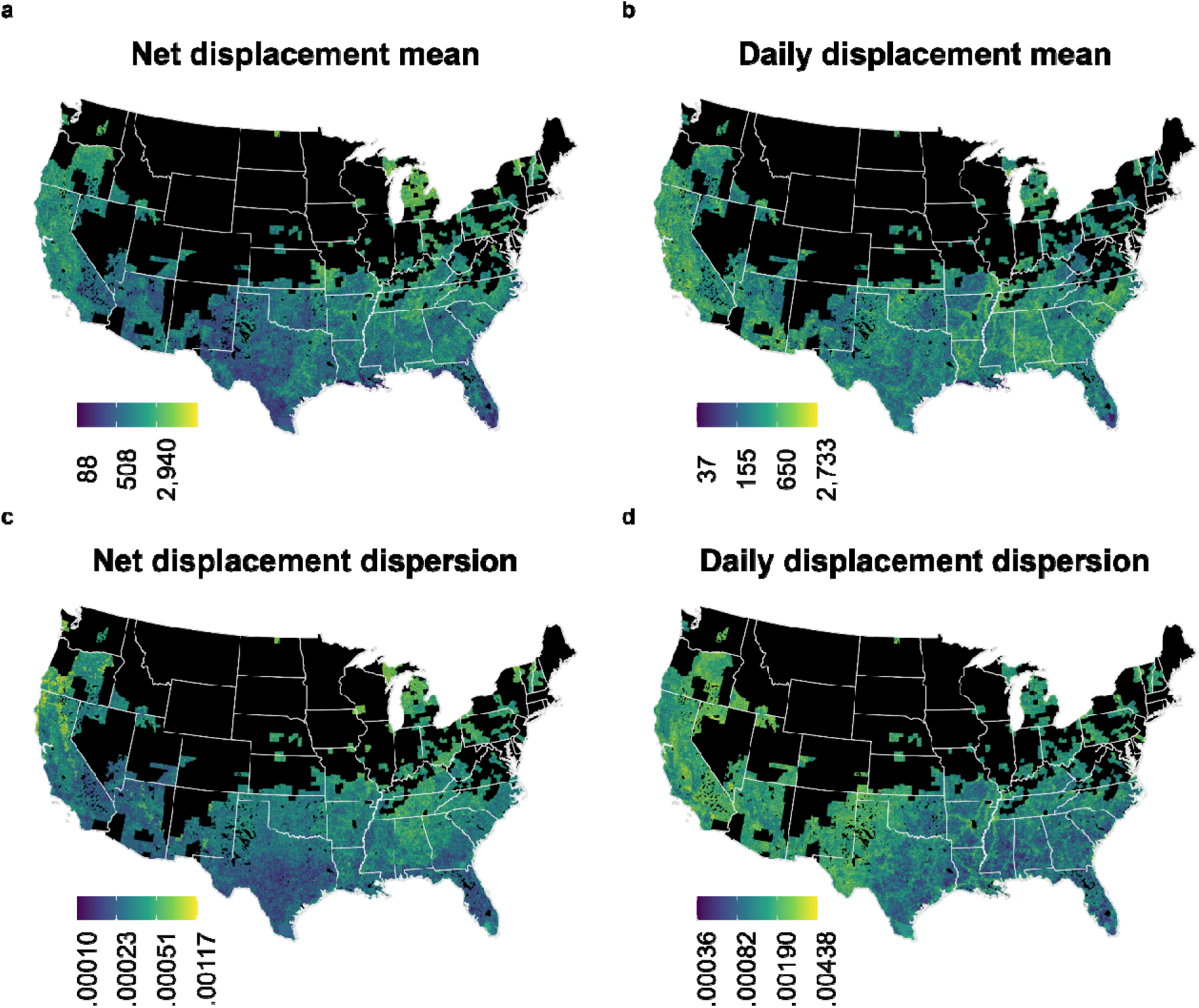
Coefficient of variation in predictions for pig displacement response variables a-d by watershed, averaged across seasons. a) daily displacement means (m); b) daily displacement dispersion; c) net displacement means (m); d) net displacement dispersion. Coefficients of variation were generated using 100 replicate predictions of the top model set of parameters and hyperparameters (determined by lowest RMSE) for each response.

**Figure S2.**
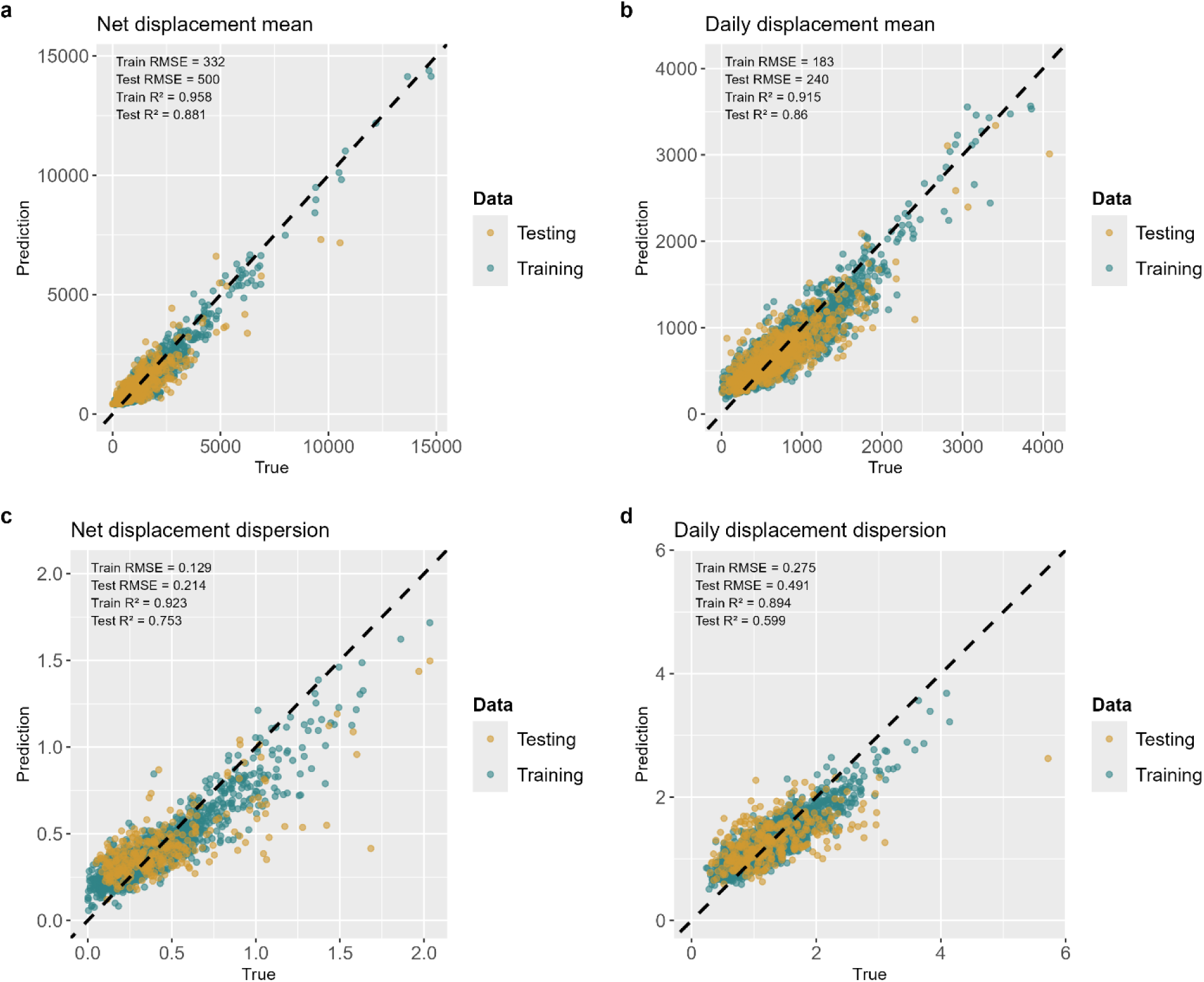
Scatterplots of observed response variables compared against predicted response variables. Testing and training data sets come from an 80:20 split of original data before training each SGBM.

**Figure S3.**
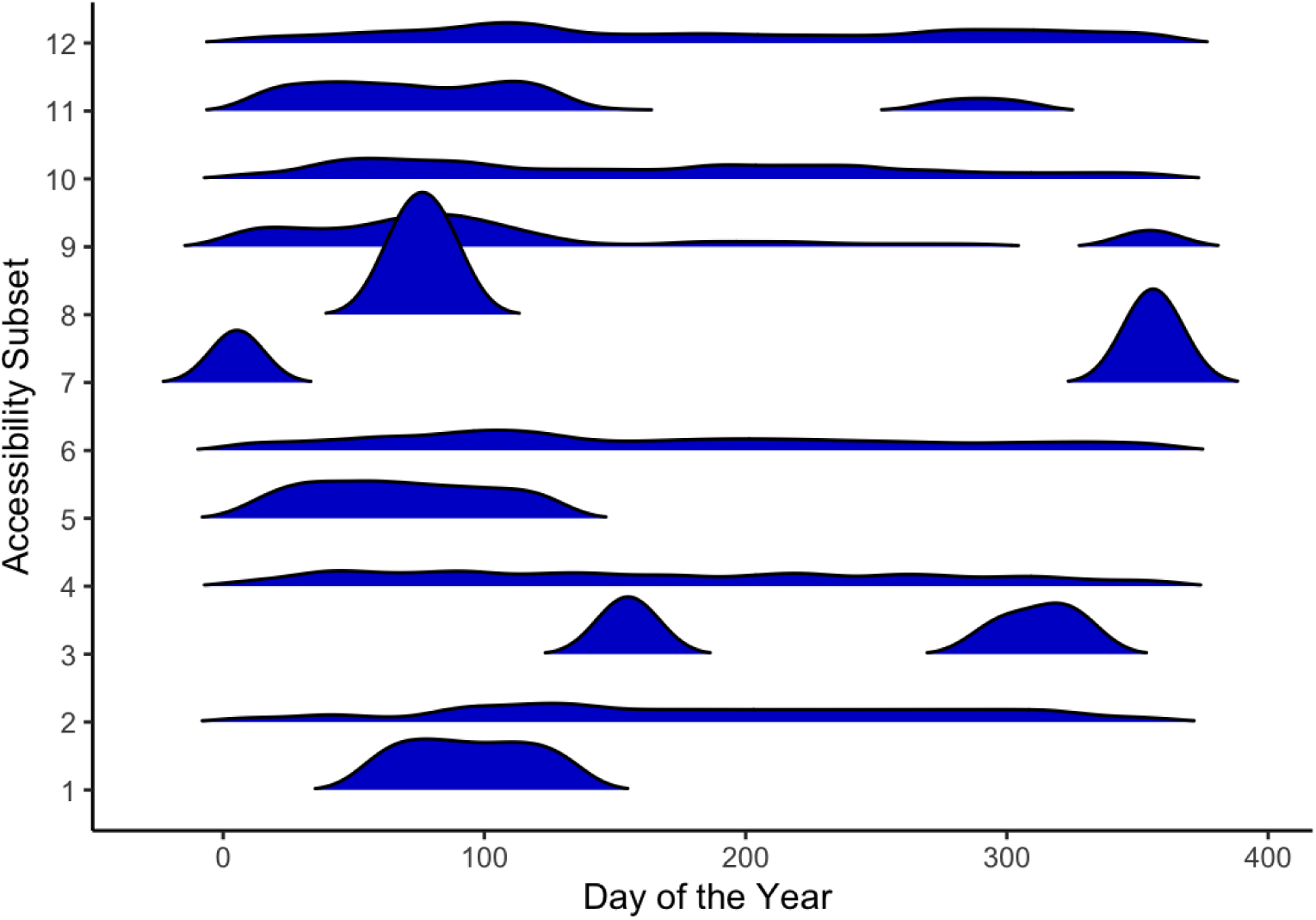
Distribution of observations over days of the year for habitat availability subsets in the step selection model, subset according to access to wetlands, urban areas, forests, and cropland

**Figure S4.**
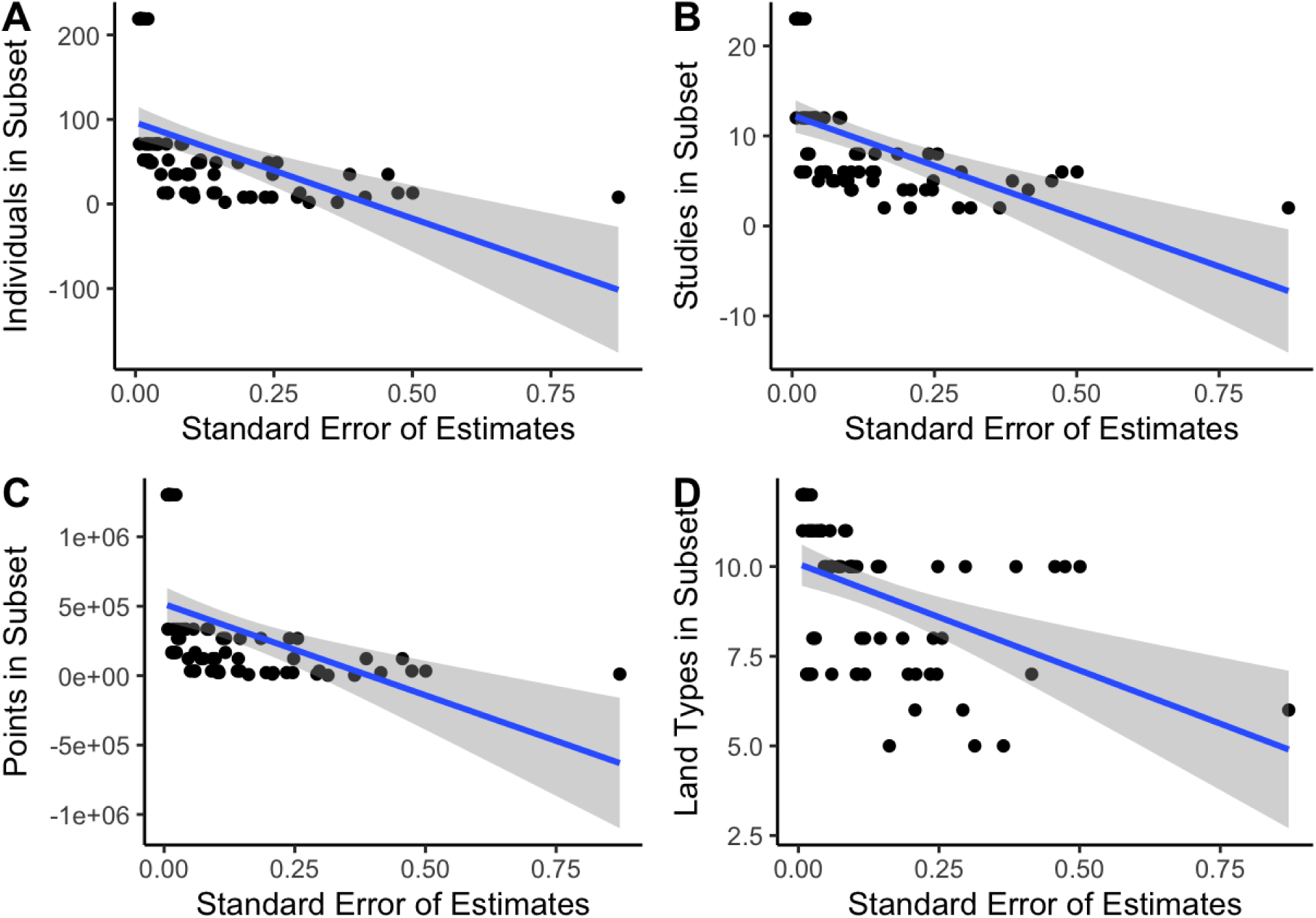
Plots of standard error of estimates for each land type from step selection model according from the leave one out analysis to A) number of pigs, B) number of studies, C) number of total GPS points and D) number of used land types in each availability subset with trend line.

**Figure S5.**
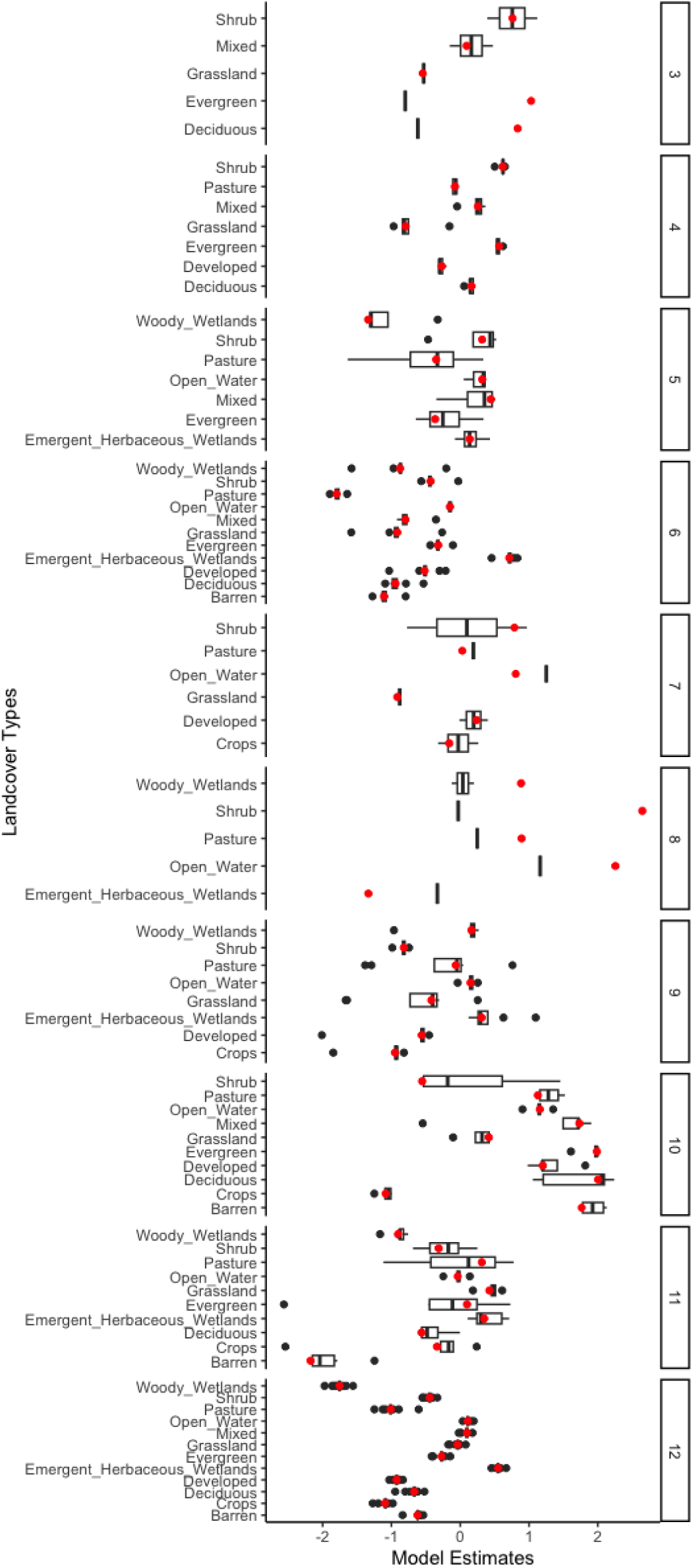
Estimates from step selection model: each panel is an accessibility subset by access to wetlands, urban areas, forests, and cropland. The black boxplot shows the estimates with one study (1-23) removed and models re-run as compared to original full model estimates (in red). Subset 1 and 2 were made of a single study and excluded from this analysis.

**Figure S6.**
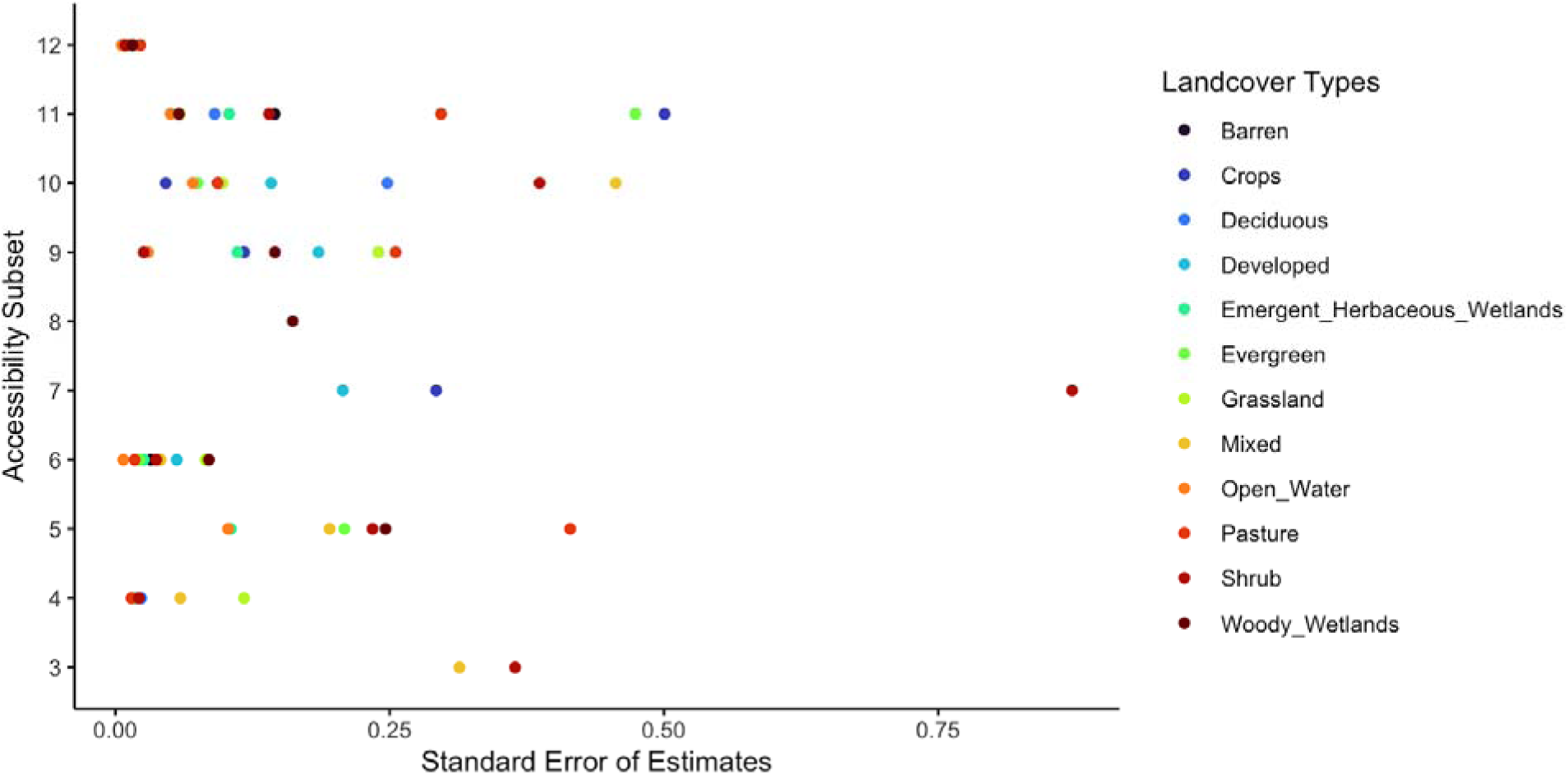
Standard error of the coefficients, subset by access to wetlands, urban areas, forests, and cropland with one study (1-23) removed and models re-run as compared to original full model estimates. Subset 1 and 2 were made of a single study and excluded from this analysis.

